# An orally available PfPKG inhibitor blocks sporozoite infection of the liver

**DOI:** 10.1101/2025.07.09.663303

**Authors:** Hanna Dhiyebi, Aminata Mbaye, Anusha Thaniana, John Gilleran, Tyler Eck, Kutub Ashraf, Karl Kudyba, Howard Fan, Steve Seibold, Kevin P. Battaile, John Siekierka, Evan Johnson, Alison Roth, Amy De Rocher, Scott Lovell, Edward B. Miller, Jacques Y. Roberge, Purnima Bhanot

## Abstract

Malaria remains a global health threat exacerbated by emerging resistance to antimalarial therapies and insecticides, climate-driven outbreaks, and limited chemoprotective options. Here, we report the characterization of **RUPB-61**, the first orally bioavailable inhibitor of *Plasmodium falciparum* cGMP-dependent protein kinase (PfPKG). **RUPB-61** prevents infection by *P. falciparum* and *P. cynomolgi* sporozoites, including the formation of hypnozoites by the latter. A single oral dose blocks liver infection by *P. berghei* sporozoites *in vivo*, demonstrating efficacy consistent with further development as a once-weekly prophylaxis based on pharmacokinetic modeling. The compound retains activity against field isolates resistant to chloroquine, mefloquine, cycloguanil, sulfadoxine and pyrimethamine, suggesting low likelihood of cross-resistance to existing antimalarials. Structural studies and free energy-based modeling guided-compound design prospectively validated the predictive accuracy of an *in silico* model of PfPKG interactions with this chemotype. While selectivity profiling identified off-target activity against human kinases, structural modeling provides a clear path for optimization. These results establish PfPKG inhibitors as promising candidates for chemoprevention and support further preclinical development of the **RUPB-61** chemotype.

**Author Summary:** Malaria remains a serious global health threat, made worse by growing resistance to existing drugs and the spread of insecticide-resistant mosquitoes. A critical gap in our prevention toolkit is the lack of safe, long-acting oral drugs that stop malaria parasites from infecting the liver - the first and obligatory step of every new infection. Here, we report the characterization of **RUPB-61**, a compound that blocks an enzyme that the malaria parasite needs to invade and survive in the liver. We show that a single oral dose completely prevents liver infection in mice, and that the compound remains active against parasites that are resistant to several currently used antimalarial drugs. Using X-ray crystallography and computer-based modeling, we determined how **RUPB-61** binds to its target in atomic detail, providing a roadmap for designing improved versions. While the compound shows some unwanted interactions with human proteins, our structural data offer clear strategies to solve this challenge. Together, these findings provide a solid foundation for developing a new class of once-weekly oral malaria prevention drugs.

## INTRODUCTION

Chemoprevention is one of two Target Product Profiles required for malaria elimination [1]. Target Candidate Profile 2 aims to address this need by developing drugs that prevent hepatic schizont and hypnozoite formation and are active against asexual stages. Such drugs will aid in reinvigorating the stalled progress in controlling malaria [2]. The first obligate step in malaria, referred to as the ‘pre-erythrocytic’ stage, is the infection by and development in the liver of *Plasmodium* sporozoites. Inhibition of sporozoite infection, through the use of vaccines or bed nets, significantly reduces disease severity and incidence [3, 4]. Drugs that target pre-erythrocytic stages are an essential component of the anti-malarial effort because even if all symptomatic malaria cases could be treated, its global eradication requires the prevention of new infections [5]. Protection is needed for vulnerable populations, like children and pregnant women, as well as populations with reduced natural immunity due to decreased exposure to parasites, such as travelers from malaria-free zones visiting endemic areas and those living in newly malaria-free areas at risk of local outbreaks [5]. Infection can be prevented through vaccines, but current vaccines are only partially effective after 4-5 doses and are recommended only for pediatric use in Africa [6, 7]. Drugs that block sporozoite infection can reduce hypnozoite formation by *P. vivax,* the most geographically widespread *Plasmodium* species, that is increasingly being detected in Africa and is the major cause of disease relapse [8, 9]. Therapies that block its liver stage maturation and/or egress will prevent relapse caused by reactivated hypnozoites and reduce the number of asymptomatic individuals who serve as reservoirs of infection and transmission [10].

Current drugs targeting pre-erythrocytic stages – primaquine, tafenoquine and atovaquone – have several drawbacks. Primaquine and tafenoquine have a common chemotype, 8-aminoquinoline, which has serious side effects. It raises the risk of hemolytic anemia in individuals with genetic deficiencies in glucose-6-phosphate dehydrogenase [11], prevalent in malaria-endemic regions [12]. In addition, primaquine is a prodrug that undergoes CYP2D6-dependent metabolism for conversion into its active metabolite [13, 14]. CYP2D6 genetic variants that are common in malaria-endemic regions are poor metabolizers of primaquine, limiting its efficacy in this population [13, 15]. Meanwhile, atovaquone is expensive, and parasites acquire resistance easily [16]. These drawbacks create an acute need for drugs that target pre-erythrocytic stages with novel mechanisms of action.

PfPKG’s function is essential in pre-erythrocytic, asexual and sexual stages of the parasite [17–19]. It is chemically and biologically validated as a drug target during the parasite’s liver infection. Its inhibition impedes the motility of sporozoites and hence their invasion of hepatocytes [20]. The development and/or exit of mature liver stage parasites from the hepatocyte in the form of merosomes is also blocked when PKG function is lost [20, 21]. Developmentally-arrested liver stages are targeted by the host immune response, and this response may potentially protect against subsequent infections [21]. PfPKG’s selective targeting is possible due to differences between its ATP-binding pocket and that of human PKG-I [22, 23].

We previously reported a structure-activity relationship (SAR) study of a trisubstituted pyrrole (**TSP**, **compound 1** [24]) with activity against PKG of related Apicomplexan parasites [25]. We used the SAR and crystallographic data to develop a free energy perturbation (FEP+) model for our series [25]. This work identified an early-lead compound, **RUPB-61** (**53** in [25]), with an IC_50_ of 13 nM against PfPKG and 50 nM against the drug-sensitive *P. falciparum* lab strain, 3D7. There is a 15-fold increase in its IC_50_ against mutant *P. falciparum* parasites carrying a Thr618Gln substitution of the ‘gatekeeper’ residue, demonstrating that its primary cellular target is PfPKG [25]. It has favorable *in vitro* pharmacokinetic (PK) properties - low preliminary cardiovascular risks as judged by hERG inhibition, high solubility and minimal first-pass metabolism, while also being more synthetically tractable due to the preferential formation of the cis isomer [25]. These data supported further investigation of the **RUPB-61** chemotype. Here, we report its *in vivo* efficacy and activity against *P. falciparum* strains resistant to current anti-malarials. We solved the structures of *P. vivax* PKG (PvPKG) bound to the compound and its enantiomer. We used these data and the SAR of closely related analogs in the series to validate the FEP+ model for PfPKG interactions with this sub-chemotype. We tested **RUPB-61**’s binding affinity for the human kinome to assess its potential for cross inhibition of human kinases. We built *in silico* models for its interactions with these kinases to identify sites for future structural modifications aimed at mitigating these liabilities. These advances will enable the rational prospective design of molecules with improved potency against PfPKG and selectivity for the parasite.

## RESULTS

### RUPB-61 prevents infection by *P. falciparum* and *P. cynomolgi* sporozoites

**RUPB-61** is active against *P. falciparum*’s asexual erythrocytic stages and *P. berghei* sporozoites infecting HepG2 cells [25]. Here, we used infection of primary human hepatocytes (PHH) to test the activity of **RUPB-61** against *P. falciparum* sporozoites. A phosphatidylinositol 4-kinase (PI(4)K) inhibitor, **KDU-691** [26] was used as a positive control. **RUPB-61** displays an IC_50_ of 79 nM in the *P. falciparum* sporozoite-PHH model with no cytotoxicity against PHH (**Figure 1A, Table 1, Supplementary Figure 1**). Next, we tested its effect on the hypnozoite-forming primate-infective species, *P. cynomolgi*. **RUPB-61** prevented the formation of hepatic hypnozoites and schizonts with an IC_50_ of 190 nM and 170 nM, respectively (**Figure 1B – C, Table 1**). These data demonstrate that the scaffold holds promise for development as pre-exposure prophylaxis against human-infective sporozoites.

**Figure 1:**
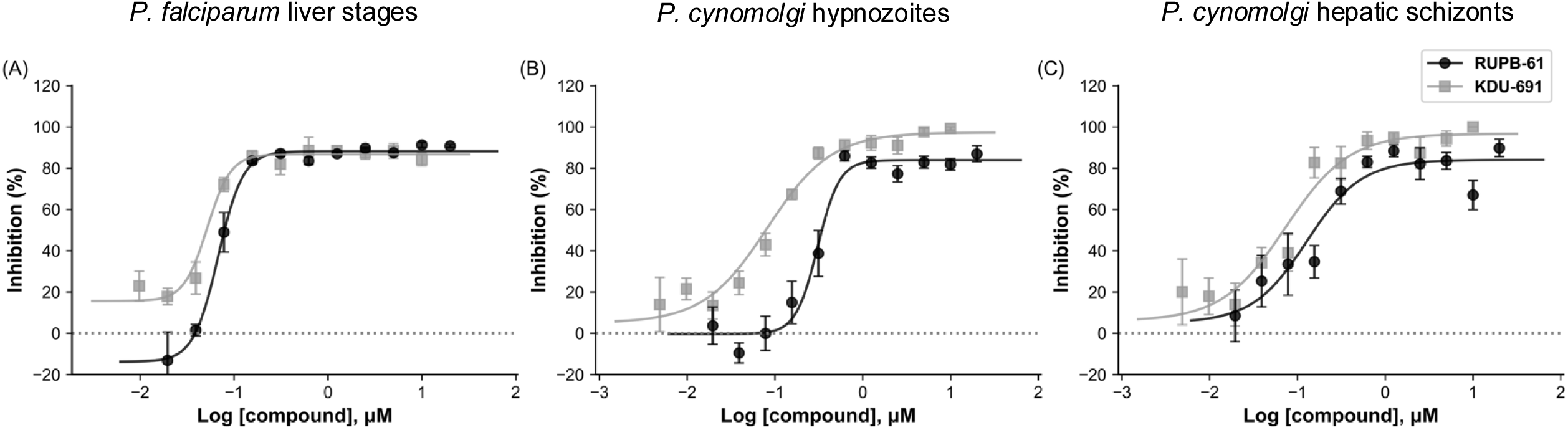
Activity of RUPB-61 in primary hepatocytes against human-infective and simian-infective sporozoites. **(A)** Effect of **RUPB-61** and **KDU-691** on *P. falciparum* infection of primary human hepatocytes, normalized to vehicle control. Infection was measured by quantifying the number of infected cells. Data shown are the mean of two biological replicates, each performed in technical duplicates. **(B)** Effect of **RUPB-61** and **KDU-691** on the number of *P. cynomolgi* hypnozoites formed in primary simian hepatocytes, normalized to vehicle control. Data shown are mean of three biological replicates, each performed in technical duplicates or triplicates. **(C)** Effect of **RUPB-61** and **KDU-691** on the number of *P. cynomolgi* schizonts in primary simian hepatocytes, normalized to vehicle control. Data shown are mean of three biological replicates, each performed in technical duplicates or triplicates.

**Table 1:** Activity of RUPB-61 against human-infective and hypnozoite-forming sporozoites. IC_50_ of **RUPB-61** and **TSP** on infection of human or simian primary hepatocytes by *P. falciparum* and *P. cynomolgi* sporozoites. **KDU-691** and tafenoquine were used as controls. Cellular cytotoxicity (CC_50_) was quantified in human and simian hepatocytes.

|  | IC<sub>50</sub> or CC<sub>50</sub> (nM) |  |  |  |
| --- | --- | --- | --- | --- |
|  | RUPB-61 | TSP | KDU-691 | Tafenoquine |
| *P. falciparum* sporozoites | 79 | na | 51 | na |
| *P. cynomolgi* hypnozoite | 190 | 400 | 110 | 85 |
| *P. cynomolgi* schizont | 170 | 320 | 42 | 41 |
| Human hepatocytes | >20,000 | na | >10,000 | na |
| Simian hepatocytes | >20,000 | 8,500 | >10,000 | 3,600 |

### RUPB-61 has a low propensity for cross-resistance

We previously reported the activity of **RUPB-61** and related stereoisomers, **RUPB-58, -59 and -60** (**51, 50** and **52**, respectively in [25]) against dihydroartemisinin (DHA)-resistant parasites that carry engineered mutations in PfKelch13 (CamC580Y) [25]. Their IC_50_ against the DHA-sensitive CamWT and DHA-resistant CamC580Y in ring-stage assays is similar [25]. Here, we assessed their cross-resistance against other clinically used antimalarial drugs by measuring their cellular activity against a panel of naturally occurring South American (7G8) and Southeast Asian (Dd2, K1 and TM90_C2B) *P. falciparum* field isolates. These strains collectively carry the most common alleles of *pfdhfr*, *pfdhps*, *pfcrt and pfmdr1* that confer resistance to chloroquine, cycloguanil, sulfadoxine, atovaquone and pyrimethamine [27]. The change in the IC_50_ of **RUPB-58** - **61** in the resistant parasites relative to 3D7 is below 5-fold. These results demonstrate that the compounds maintain their activity against multidrug resistant *P. falciparum* parasites (**Table 2, Supplementary Figure 2**).

**Table 2:** PfPKG inhibitors are active against multi-drug resistant *P. falciparum*. IC_50_ of **RUPB-58** - **61** against *P. falciparum* isolates resistant to current antimalarial drugs. Data shown are mean (standard error) of 3 biological replicates.

|  | **3D7**<br>**(nM)** | **Dd2-B2**<br>**(nM)** | **TM90_C2B**<br>**(nM)** | **K1**<br>**(nM)** | **7G8**<br>**(nM)** |
| --- | --- | --- | --- | --- | --- |
| **RUPB-58** | 170<br>(30) | 467<br>(113) | 122<br>(24) | 239<br>(47) | 57<br>(42) |
| **RUPB-59** | 80<br>(23) | 353<br>(45) | 120<br>(18) | 148<br>(1) | 135<br>(25) |
| **RUPB-60** | 50<br>(53) | 144<br>(3) | 80<br>(44) | 81<br>(1) | 28<br>(9) |
| **RUPB-61** | 50<br>(32) | 129<br>(10) | 139<br>(54) | 42<br>(0.3) | 29<br>(6) |
| **DHA** | 2<br>(0.3) | 3<br>(1) | ND | 2<br>(0.01) | 0.5<br>(0.02) |

Replacement of the PfPKG ‘gatekeeper’ Thr618 with larger amino acids (e.g. Met, Phe, Tyr, Leu, Ile or Gln) are expected to lead to resistance to **RUPB-61** and structurally related analogs since their bulkier sidechains decrease the volume of the binding pocket. Previous *in vitro* resistance studies using PfPKG inhibitors of different chemotypes either failed to select resistant mutants or found only low-level resistance from mutations in non-targets [28, 29]. Here, we evaluated the risk of naturally occurring escape ‘gatekeeper’ mutants. We examined high-resolution genetic information from 20,892 parasite samples present in the MalariaGEN Pf7 database. These include isolates from major malaria-endemic regions of the world, laboratory clonal and mixed samples, and the results of experimental genetic crosses. Analysis revealed natural variations in the PfPKG protein – 493 synonymous changes (in 4823 samples) and 523 non-synonymous changes (in 1164 samples) (**Supplementary Table 1**). At the ‘gatekeeper’ position, two isolates carried synonymous changes (Thr618Thr), and one isolate carried a non-synonymous mutation (Thr618Ser). The number of samples with changes in the ‘gatekeeper’ position was not statistically significant compared to the total samples analyzed. The extreme rarity of non-synonymous mutations at the ‘gatekeeper’ position suggests that randomly occurring pre-existing mutations are unlikely to facilitate resistance to the **RUPB-61** chemotype through this mechanism under natural conditions.

### RUPB-61 is orally available

One of the requirements for a chemoprotective agent is oral administration at a frequency of at most once-weekly and ideally, once-monthly. Achieving these benchmarks requires compounds with extended oral half-lives. We tested **RUPB-61**’s ADME properties in mice after oral and intravenous dosing at 10 mg/kg. Plasma concentrations from both dosing regimens were identical from 2 - 24 h (**Figure 2**). The calculated oral bioavailability is 91%, and the oral half-life is approximately 9 h (**Table 3, Supplementary Table 2**). Allometric scaling translates this to approximately 2-4 days in humans [30]. The compound maintains exposure at >100 nM (∼2x EC_50_^3D7^) for 24 h with relative clearance around 50 mL/min/kg and large volume of distribution of 47.3 L/kg (**Figure 2, Table 3, Supplementary Table 2**).

**Figure 2:**
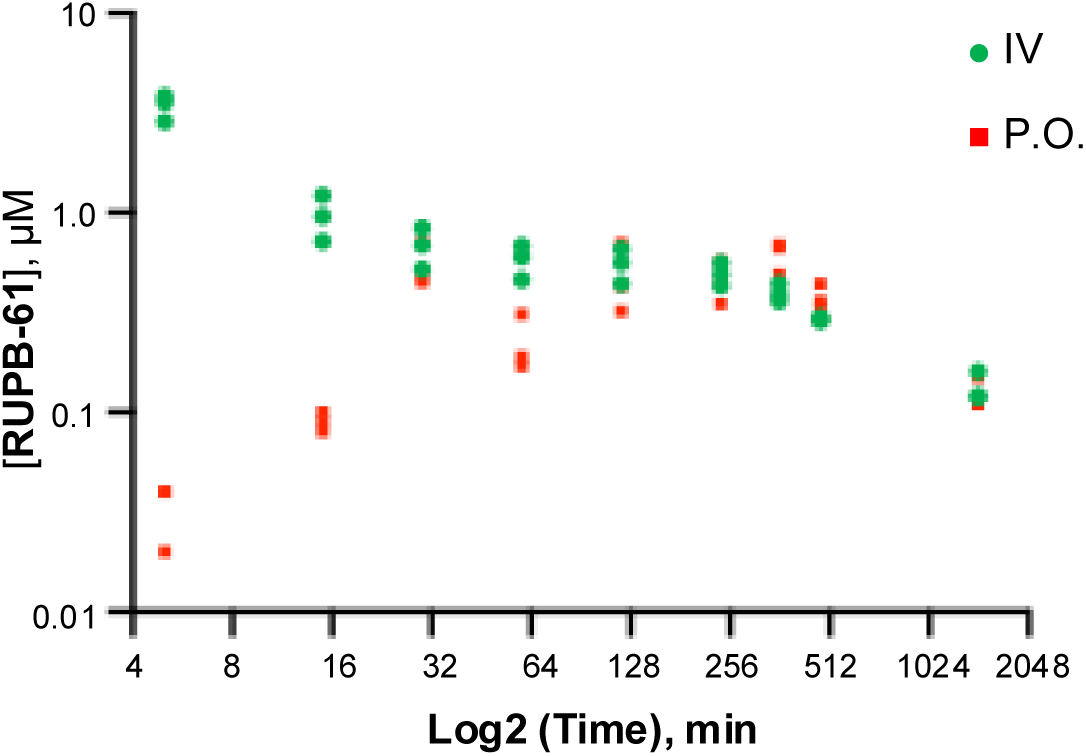
Plasma half-life of RUPB-61 *in vivo*. Plasma concentration in mice was measured over a 24 h period after intravenous (IV) and oral (PO) dosing with 10 mg/kg (n = 3 per route).

**Table 3:** *In vivo* assessment of RUPB-61 ADME properties. Pharmacokinetic properties were determined after oral (PO) or intravenous (IV) administration of compound to mice. Data shown are average of 3 mice. T_max_: time to maximum distribution; C_max_: Maximum serum concentration; AUC: Area under the Curve; Cl: clearance; INF: infinite; MRT: Mean Residence Time; V_ss_: Volume of distribution

| **PO** | **T<sub>1/2</sub>**<br>(h) | **T<sub>max</sub>**<br>(h) | **C<sub>max</sub>**<br>(μM) | **AUC<sub>last</sub>**<br>(μM*h) | **AUC<sub>INF_obs</sub>**<br>(μM*h) | **AUC<sub>Extrap_obs</sub>**<br>(%) | **Cl<sub>obs</sub>**<br>(ml/min/kg) | **Bioavailability**<br>%F |  |
| --- | --- | --- | --- | --- | --- | --- | --- | --- | --- |
| Average | 8.71 | 4.17 | 0.63 | 7.29 | 8.9 | 18.16 | 54.31 | 91% |  |
| **IV** | **T<sub>1/2</sub>**<br>(h) | **T<sub>max</sub>**<br>(h) | **C<sub>max</sub>**<br>(μM) | **AUC<sub>last</sub>**<br>(μM*h) | **AUC<sub>INF_obs</sub>**<br>(μM*h) | **AUC<sub>Extrap_obs</sub>**<br>(%) | **Cl<sub>obs</sub>**<br>(ml/min/kg) | **MRT<sub>INF_obs</sub>**<br>h | **Vss<sub>obs</sub>**<br>L/kg |
| Average | 11.78 | 0.08 | 3.42 | 7.43 | 9.8 | 23.60 | 49.25 | 16.27 | 47.29 |

### A single oral dose of RUPB-61 is efficacious in a murine model of liver infection

The similarity in **RUPB-61**’s activity against *P. falciparum* and *P. berghei* sporozoites strongly supports the use of *P. berghei* as a model for testing compound activity *in vivo*. We tested the efficacy of **RUPB-61** and the parent compound, **TSP**, against liver infection by *P. berghei* sporozoites expressing luciferase and GFP (PbLuc-GFP). In the first experiment, mice were administered three doses, intravenously (IV) or orally (PO), of compound (10 mg/kg) - two prior to sporozoite inoculation to block sporozoite invasion of hepatocytes and the third at 36 hpi to prevent egress of liver stages (**Figure 3A**). *In vivo* bioluminescent imaging (IVIS) of mice was utilized to detect and quantify relative parasitemia in the liver and flow cytometry was used to detect the appearance of blood stage parasites. Vehicle-treated mice were used as a negative control. For both treatments, liver parasitemia was detectable at 44 h post infection (hpi) in all vehicle-treated mice (10/10) (**Figure 3B-C, Supplementary Figure 3**). Additionally, a majority of mice treated (IV) with **TSP** (3/5) displayed liver infection at 44 hpi. In contrast, none of the mice treated with **RUPB-61** (10/10) showed liver parasitemia as late as 144 hpi (**Figure 3B-C, Supplementary Figure 3**).

**Figure 3:**
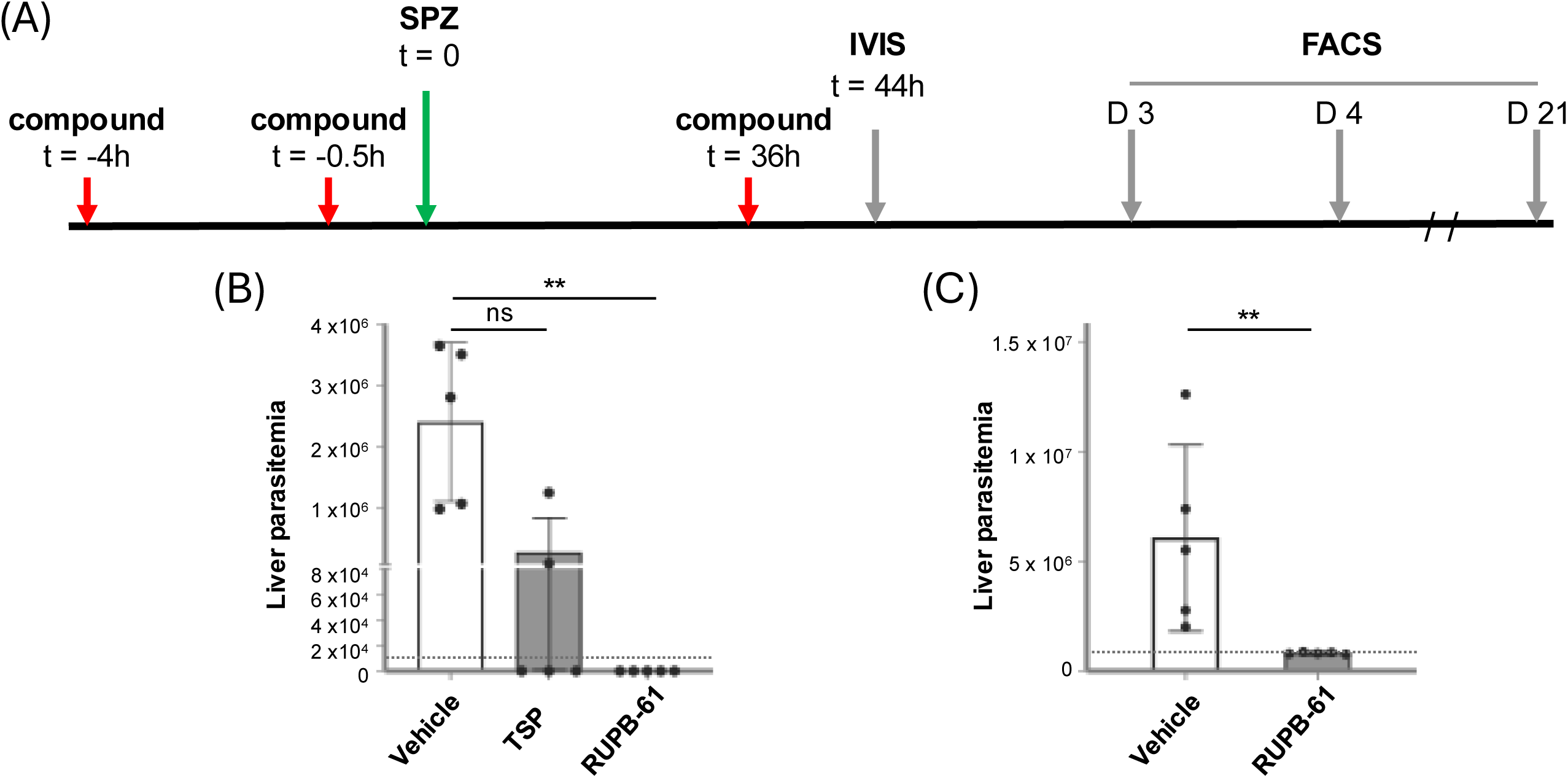
Efficacy of RUPB-61 *in vivo*. **(A**) Experimental design to test the efficacy of **RUPB-61** against PbLuc-GFP sporozoite-initiated infection in mice. Mice were administered 3 doses of compound intravenously or orally. Liver parasitemia was quantified using bioluminescent imaging and blood stage parasitemia was detected using FACS. **(B)** Liver parasitemia at 44 hpi in infected mice administered compounds IV. **(C)** Liver parasitemia at 44 hpi in infected mice administered compounds orally. Parasitemia was measured in radiance (p/sec/cm^2^/sr). Dotted line represents background signal.

A decrease in liver stage parasitemia is expected to delay the appearance of blood stage parasites, as there are fewer hepatic merozoites to initiate erythrocytic infection. Therefore, we quantified the time taken to detect blood stage parasites after sporozoite inoculation (pre-patent period) in these mice. The pre-patent period in vehicle-treated animals was 3.6 days under IV administration and 4.6 days in oral treatment, while none of the **RUPB-61**-treated mice displayed blood stage parasites in the 3 week observation period (**Table 4**). In the group that received 3 intravenous doses of **TSP**, 2/5 mice did not show blood stage parasites for 3 weeks pi. This implies that the lack of detectable liver stage parasitemia at 44 hpi represents a complete block in sporozoite infection. In the remaining **TSP**-treated mice that had detectable liver parasitemia at 44 hpi, the pre-patent period trended longer than vehicle-treated mice of the same cohort (4.3 days versus 3.6 days) (**Table 4**), although the difference was not statistically significant.

**Table 4:**
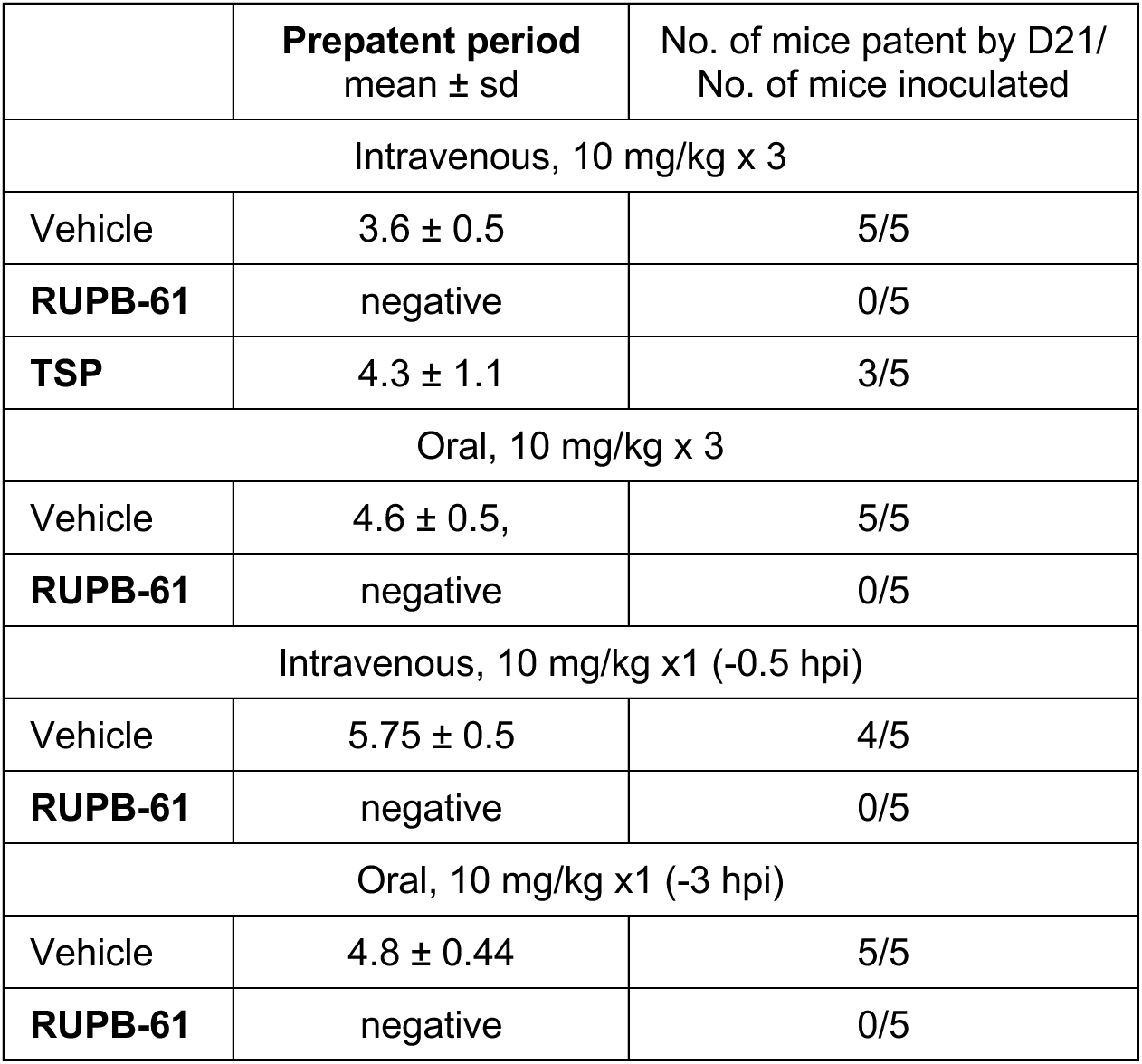
Effect on RUPB-61 on the prepatent period of infection. Prepatent period of sporozoite infection was determined through detection of PbLuc-GFP asexual blood stages using flow cytometry. sd: standard deviation

|  | **Prepatent period**<br>mean ± sd | No. of mice patent by D21/<br>No. of mice inoculated |
| --- | --- | --- |
| Intravenous, 10 mg/kg x 3 |  |  |
| Vehicle | 3.6 ± 0.5 | 5/5 |
| **RUPB-61** | negative | 0/5 |
| **TSP** | 4.3 ± 1.1 | 3/5 |
| Oral, 10 mg/kg x 3 |  |  |
| Vehicle | 4.6 ± 0.5, | 5/5 |
| **RUPB-61** | negative | 0/5 |
| Intravenous, 10 mg/kg x1 (-0.5 hpi) |  |  |
| Vehicle | 5.75 ± 0.5 | 4/5 |
| **RUPB-61** | negative | 0/5 |
| Oral, 10 mg/kg x1 (-3 hpi) |  |  |
| Vehicle | 4.8 ± 0.44 | 5/5 |
| **RUPB-61** | negative | 0/5 |

Finally, we tested the efficacy of single-dose **RUPB-61** when administered intravenously or through oral gavage. Based on the *in vivo* pharmacokinetic data (**Figure 2**), the oral dose was provided at -3 hpi and the intravenous dose at -0.5 hpi to ensure high levels of compound in plasma at the time of sporozoite inoculation. Vehicle-treated mice (9/10) were positive for blood stage parasites by 4.8 days and 5.75 days, respectively whereas none of the **RUPB-61**-treated mice (10/10) developed detectable blood stage infection for the 3-week monitoring period (**Table 4**). We conclude that **RUPB-61** is sufficient to prevent infection of the liver by *P. berghei* sporozoites at a single, dose of 10 mg/kg delivered orally.

### RUPB-61 has moderate selectivity for human kinases

A major issue affecting the further development of **RUPB-61** is its human-kinome selectivity. To identify its significant off-target liabilities, we quantified its binding affinity for 400 unique human kinases (**Supplementary Figure 4**). The screen consists of binding competitions between the test compound and control ligands for each of the target human kinases. At 1 μM **RUPB-61** displaced >99% of the control ligands for CIT, ROCK1 and ROCK2, hence displaying moderate selectivity against the human kinome. We then determined the binding affinity (Kd) of **RUPB-61** for the three kinases (**Table 5**). The Kd for CIT was the lowest, identifying it as the foremost off-target of **RUPB-61**.

**Table 5:**
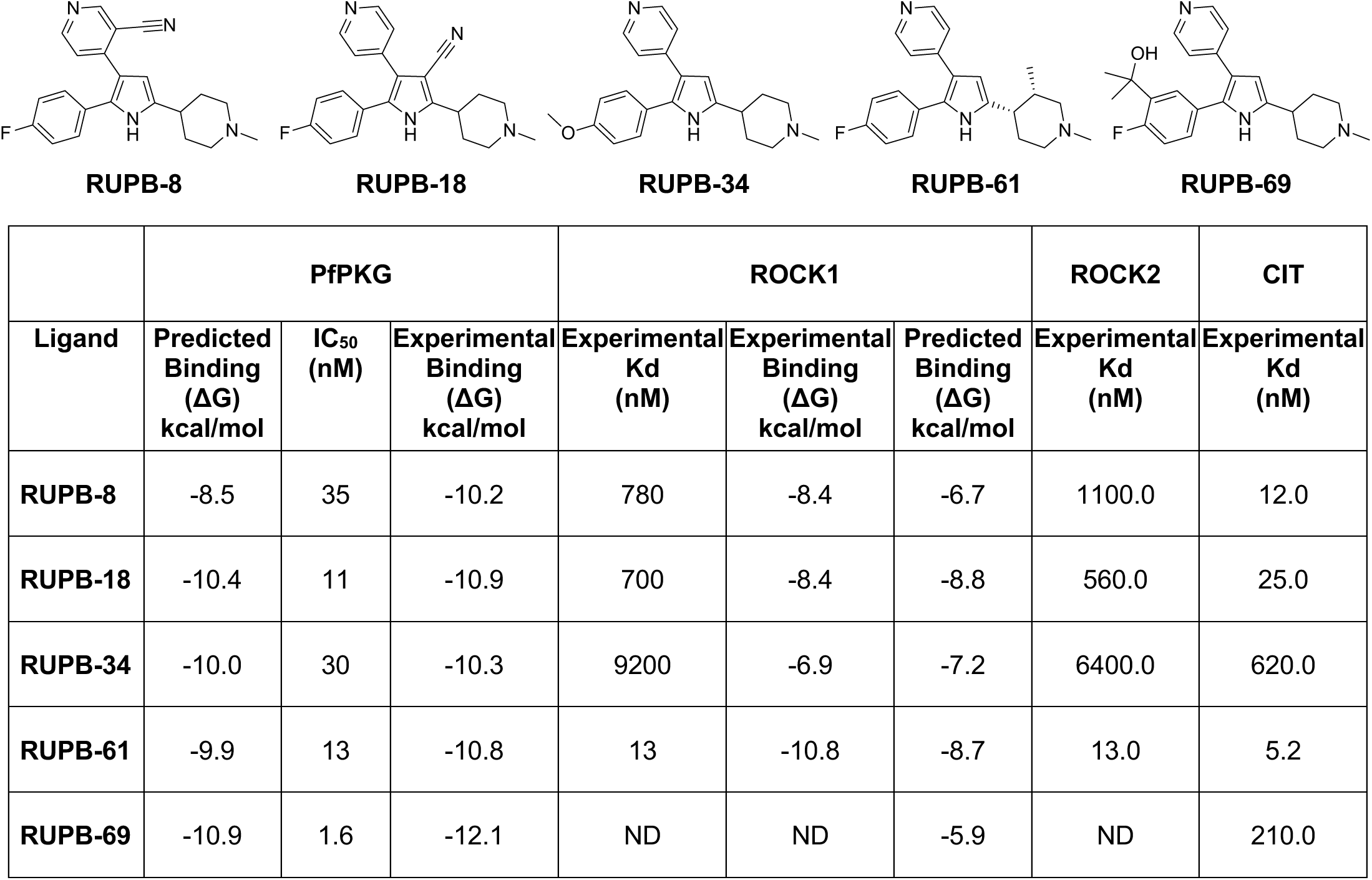
Initial SAR investigation of the RUPB-61 series for human kinase off-targets. Binding affinities (Kd, nM) of compounds for CIT, ROCK1 and ROCK2 were determined experimentally and predicted using a toxicology tool. Experimental ΔG was calculated using the relationships IC_50_ ≈ Kd and ΔG = -RTln(K_d_) with 300K for temperature. ND: not determined

Further development of **RUPB-61** will require de-risking these kinases. In order to better understand their interaction with the **RUPB-61** series of compounds, we determined the binding affinities of analogs with distinct structural modifications, **RUPB-8, -18, -34** and **-69** (**Table 5**), against ROCK1/2 and CIT. The compounds were selected because they carry significant structural modifications relative to **RUPB-61** and maintain activity against PfPKG (**Table 5**). Compared to **RUPB-61,** the Kds of **RUPB-8** (3-cyanopyridine), **RUPB-18** (3-cyanopyrrole) and **RUPB-34** (4-methoxyphenyl) are 40 - 700-fold higher for ROCK1 and ROCK2. The Kds of **RUPB-18**,**-34 and -69** (dimethylbenzyl alcohol) for CIT were 5 - 125-fold higher compared to **RUPB-61**. In addition to experimental SAR, we utilized a predictive toxicology tool (Schrödinger Predictive Toxicology Models) for computational assessments of relative binding affinities of a given compound for human kinases. CIT was not available within the *in silico* kinase panel. However, since ROCK1 and CIT are both members of the AGC kinase family we postulated that ROCK1 modeling may also serve as a proxy for CIT and ROCK2. We compared experimental and model-predicted Kds (kcal/mol) of ROCK1 for **RUPB-8, -18, -34, -61** and **-69** (**Table 5**). The Pearson correlation coefficient squared (R^2^) for their measured and predicted ROCK1 ΔG is a 0.34. This contrasts with an R^2^ of 0.55 for our predicted IFD-MD PfPKG model (**Table 6**). The modest R^2^ for the ROCK1 model is most likely a reflection of the limited size of the data set. With additional data points and further refinement, the ROCK1 model should discriminate between weak, moderate and strong binders, leading to more accurate predictions.

**Table 6:** Comparison of the experimental PvPKG model. The overall FEP statistics of the two models in terms of the squared Pearson correlation coefficient R^2^, between predicted and experimental ΔG and the root-mean-squared error (RMSE) of all pairwise ΔΔGs. In parentheses are the 95% confidence interval RMSE values using bootstrapping.

| Model | FEP R<sup>2</sup> | FEP RMSE (kcal/mol) |
| --- | --- | --- |
| PvPKG Experimental Structure | 0.57 | 1.29 [1.23, 1.36] |
| PfPKG IFD-MD Model | 0.55 | 1.32 [1.26, 1.39] |

### Atomic structure of PvPKG bound to RUPB-61 and its enantiomer, RUPB-60

Further development of **RUPB-61** will include improvement in its potency. Rational design of analogs will be facilitated by structural information. Towards this end, we solved the structure of **RUPB-61** bound to *P. vivax* PKG (PvPKG) at 2.8 Å (PDB 9P74) (**Figure 4, Supplementary Table 3**). We compared the predicted **RUPB-61**-bound PfPKG structure from our prior work [25] with the experimental **RUPB-61**-bound PvPKG structure. These results showed that ligand poses are extremely similar. Structural superposition of the two kinase structures leads to a ligand heavy-atom RMSD of 0.82 Å. In both structures, the hinge interaction is formed using the pyridine nitrogen as an acceptor for a backbone NH (Val621 in PfPKG, Val614 in PvPKG) with aromatic CH interactions ortho to the pyridine nitrogen interacting with backbone carbonyls (Val621 and Glu619 in PfPKG, only Glu612 in PvPKG). The most significant difference between the two models lies in the conformation of the ‘P-loop’. In the predicted PfPKG structure, the P-loop dips down towards the ligand and in doing so allows Asp682 of the DFG motif (Asp-Phe-Gly motif) to hydrogen bond with the Thr551 side chain hydroxyl and the backbone NH in the P-loop. This Asp forms a hydrogen bond to the ligand pyrrole NH. In the experimental PvPKG structure, the P-loop is pointing up and away from the ATP pocket. The homologous Thr in PvPKG, Thr544 is unable to form an interaction with the DFG motif Asp, Asp675 in PvPKG, and evidently the hydrogen bond with the pyrrole is not formed. An additional difference is in the interaction with Glu625 in PfPKG. In the PvPKG structure, the equivalent Glu618 is forming a salt bridge to the ligand piperidine while in the PfPKG structure, Glu625 points out into solution.

**Figure 4:**
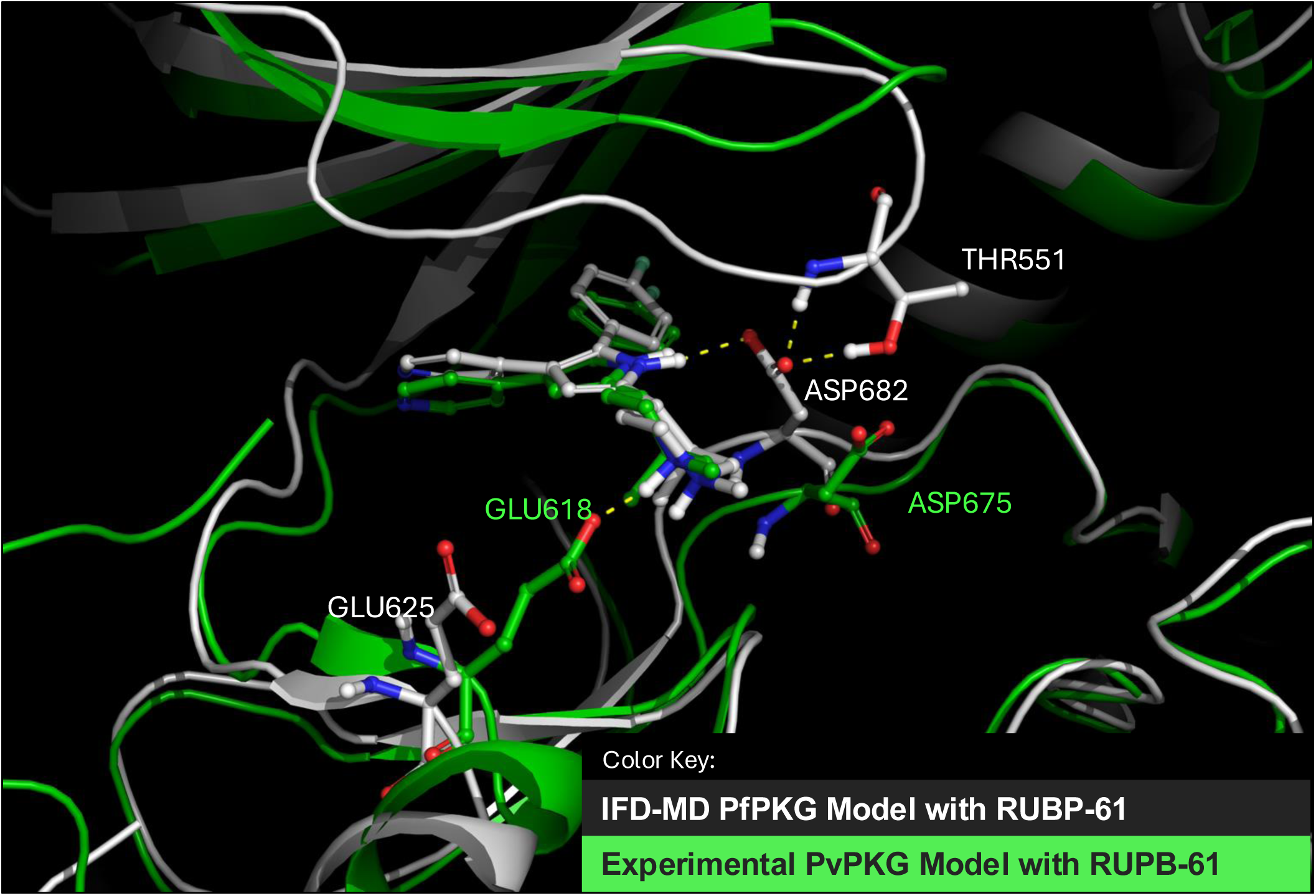
Comparison of the predicted PfPKG structure with the experimental PvPKG structure for RUPB-61. Overlay of the predicted PfPKG structure with **RUPB-61** [25] (white) and the experimental PvPKG structure (green). The two most significant differences between the two models are in the interactions of two carboxylates, E625 (E618 in PvPKG) which forms a salt-bridge to the ligand piperidine only in PvPKG and D682 (D675 in PvPKG) which only forms a charged hydrogen bind to the pyrrole NH in the PfPKG model.

**RUPB-61** is chiral at the 3- and 4-positions of the piperidine ring. To understand the effect of chirality on its interactions with PfPKG, the structure of its diastereomer, **RUPB-60,** was also solved in PvPKG (PDB 9P75). **RUPB-60** places a methyl substituent towards the solvent-exposed side of the ATP binding pocket (**Supplementary Figure 5, Supplementary Table 3**). In the **RUPB-60** structure, the C-terminal loop is resolved with an additional three residues compared to the **RUPB-61** structure. These additional three residues include Ala816 which hydrophobically packs against the methyl piperidine substituent. This Ala is conserved in PfPKG as Ala823, however, the additional hydrophobic packing does not appear to affect improved potency as **RUPB-60** and **RUPB-61** have similar IC_50_ against PfPKG variants [25].

To address whether the differences between the IFD-MD PfPKG model and experimental PvPKG model are material to modeling, we performed FEP using an identical congeneric series of compounds. We selected our previously reported compounds [25] as well as four additional compounds (one of which is represented as a pair of stereoisomers) which were prospectively designed using our prior IFD-MD model.

The statistics of the two models appear quite similar with a marginal improvement using the experimental PvPKG model (**Table 6, Supplementary Figure 6**). The RMSE values have overlapping 95% confidence intervals which is consistent with the two models appearing near equivalent in FEP. This would suggest that the differences between the two models may be due to thermal fluctuations, for example the solvent-exposed piperidine salt-bridge, as the crystal structure is solved within a 100K liquid nitrogen stream while the IFD-MD model is simulated at 300K. This also suggests that the PvPKG model and X-ray structures are valid for use as a proxy for PfPKG during molecule design.

### Experimental confirmation of FEP+ model

The solved structures of PvPKG bound to either **RUPB-61** or **RUPB-60** match qualitatively and quantitatively (**Table 6, Supplementary Table 3**) to our previously reported FEP+ model of PfPKG [25]. For orthogonal validation of the FEP+ model, we designed molecules to test some key predictions (**Table 7**). **RUPB-68, RUPB-69**, and **RUPB-72** explore displacement of a collection of high energy waters identified using WaterMap in our prior publication [25] and replacement of these interactions with an added hydroxyl or ester group compared to **TSP**. As a negative control, we tested the model’s accuracy at predicting weaker compounds by replacing the methyl groups in **RUPB-69** with ethyl groups in **RUPB-73.** These groups are expected to clash in the binding cavity, resulting in weaker activity as confirmed by experimental measurements (**Table 7**). These data are included in the aggregate statistics shown in **Table 6** with individual ligand statistics in **Supplementary Table 4**, but the predicted versus experimental affinity for these prospective ligands are shown separately in **Table 7**. Errors between prediction and experiment are around 1 kcal/mol which is consistent with published FEP+ performance using high quality crystal structures [31]. Both calculations (FEP+ in PvPKG and PfPKG) are run at 25 ns, 5x larger than the default, to help eliminate any random errors due to insufficient sampling or convergence. The congruence between these data, the co-crystal structures and predictions of the FEP+ model demonstrates the validity of the FEP+ model. This supports the use of the FEP+ model for future prospective design of more potent PfPKG inhibitors and for maintaining potency while eliminating undesirable human kinase binding.

**Table 7:** Predicted and experimentally determined prospective inhibition of PfPKG. IC_50_ was converted from concentration to kcal/mol using the relationships IC_50_ ≈ Kd and ΔG = -RTln(K_d_) with 300K for the temperature.

|  | **PfPKG IC<sub>50</sub>**<br>**(nM)** | **PfPKG (ΔG)**<br>**(kcal/mol)** | **FEP+-predicted (ΔG)**<br>**(kcal/mol)** |
| --- | --- | --- | --- |
| **RUPB-68** | 34.5 (6.4) | -10.3 | -10.9 |
| **RUPB-69** | 1.6 (0.6) | -12.1 | -10.9 |
| **RUPB-72** | 17.2 (0.4) | -10.7 | -11.1 |
| **RUPB-73** | 256.5 (174.6) | -9.1 | -8.1 |

The IFD-MD model can also be used to predict the binding affinity of the Thr618Ser mutation found in one isolate in MalariaGen database. We used the protein FEP approach, which assumes the ligand remains roughly unchanged while introducing the T618S mutation *in silico* to PfPKG. We measured the change in binding free energy (ΔΔG) from this mutation. FEP modeling predicts that mutating T618 to S618 improves the binding affinity to the ligand slightly by 0.5 ± 0.43 kcal/mol, indicating that this mutation is not predicted to significantly affect binding. These calculations strongly suggest that the parasites carrying the Thr618Ser mutation in PfPKG will remain sensitive to **RUPB-61**.

## DISCUSSION

The global malaria landscape is under mounting threat due to resistance to artemisinin-based combination therapies and the spread of insecticide-resistant mosquito vectors. Climate-driven instability further compounds the risk, exemplified by the 5-fold surge in malaria cases in Pakistan during the 2022 floods. Even historically malaria-free regions, such as the United States, reported locally acquired infections in 2023 - the first in two decades - highlighting the urgent need for sustained innovation in prevention strategies.

A cornerstone of renewed malaria control efforts will be the development of drugs targeting pre-erythrocytic stages. These agents can complement artemisinin-based therapies in three-drug combinations or synergize with next-generation vaccines to provide robust chemoprotection, particularly for seasonal transmission or endgame elimination. Protection is needed for pregnant women, children, individuals traveling from malaria-free zones into endemic areas and for populations in newly malaria-free areas at risk of local malaria outbreaks, as these groups have reduced natural immunity to infection and disease [5]. However, there is a significant dearth of candidates that meet the stringent potency, pharmacokinetic, and safety criteria required for long-acting prophylactic agents, especially those intended for monthly oral or quarterly parenteral administration.

Our study identifies **RUPB-61** as the first orally bioavailable inhibitor of PfPKG with demonstrated single-dose efficacy in a murine model. PfPKG is a compelling target for chemoprotection, given its dual essential functions in liver stage and blood stage egress and invasion. The lower parasite burden and lack of replication cycles in the liver further increase the vulnerability of the pre-erythrocytic stages to kinase inhibition. **RUPB-61** exploits two temporally distinct windows of parasite susceptibility - during hepatocyte invasion and again during merosome formation [20, 21] - positioning it as a versatile candidate for both prevention and treatment. The inhibitory effect of PfPKG inhibitors on *P. cynomolgi* sporozoites suggests that, in addition to preventing or mitigating disease, they could reduce the formation of *P. vivax* hypnozoites, which are the major cause of vivax-malaria relapse and the number of asymptomatic individuals who serve as reservoirs of *P. vivax* infection and transmission [10]. Studies in FRG huHEP mice will further define its efficacy against *P. falciparum* sporozoites and potential activity against *P. vivax* hypnozoite formation [32].

Importantly, PfPKG inhibitors fulfill the chemoprevention TPP’s requirement of a mechanism of action distinct from current antimalarials. Our cross-resistance studies demonstrate that **RUPB-61** maintains potency in blood stage growth assays against parasites resistant to chloroquine, cycloguanil, sulfadoxine, pyrimethamine or dihydroartemisinin [25]. These findings suggest potential for co-formulation with existing frontline therapies without antagonism or cross-resistance, but this needs to be tested in combination drug assays.

Resistance to PfPKG inhibitors appears infrequent and biologically costly. Laboratory studies and data from gatekeeper mutants (e.g. T618Q) suggest that resistance-conferring mutations compromise ATP binding and enzymatic efficiency [33, 34], reducing sporozoite yield and infectivity [20]. This supports the evolutionary robustness of the target and a low likelihood of transmission of strains arising from gatekeeper mutations. Future studies in *P. falciparum* mosquito transmission models will be crucial for validating this prediction. In addition, studies on lab-evolved resistance under drug pressure are needed to estimate the minimum inoculum of resistance and the susceptibility of the scaffold to the development of resistance in parasites.

The pharmacokinetic profile of **RUPB-61** is also promising. Its similar IV and oral plasma concentrations - unusual for kinase inhibitors - result from high bioavailability and long oral half-life, which are key parameters for preclinical candidate advancement. The long half-life of the compound can result from intrinsic metabolic stability and/or depoting in tissues. **RUPB-61**’s volume of distribution suggests the latter. Since **RUPB-61** is not highly lipophilic, its tissue depoting is unlikely to result from non-specific binding to membranes. We speculate that it may result from active uptake by transporters into tissues.

Structure-guided design has been instrumental in advancing the **RUPB-61** chemotype. Prospective compounds synthesized using the IFD-MD model and FEP+ affinity predictions, prior to any retrospective fitting, validated the predictive power of the model’s. The ability to design novel, high-affinity ligands *de novo* represents a critical capability for advancing preclinical optimization and derisking development. One liability of the current scaffold is off-target activity against human kinases, notably CIT and ROCK1/2. However, our structure-based modeling provides a rational framework for selectivity optimization. Our results identify the fluorophenyl ring as an attractive target for substitutions aimed at improving selectivity. Importantly, these modifications can significantly enhance potency against PfPKG, for example the 10-fold improvement in the IC_50_ of **RUPB-69** compared to **RUPB-61**. Future work will develop and apply experimental SAR across a larger number of analogs to drive iterative molecule design. These SAR data will also be used to build a more accurate IFD-MD model of ROCK1 (with the ROCK1 model serving as a proxy for the related ROCK2 and CIT). Greater accuracy should lead to predicted binding affinities that are within 1 kcal/mol of experimental determinations. With this refinement, the *in silico* models can be used to triage prospective molecules based on selectivity for the human kinome, hence assisting in rationalizing and prioritizing compound synthesis. Previous studies demonstrate that optimization of *in silico* models can be achieved using a ligand set of 9 – 10 compounds comprising strong, moderate and weak binders [35].

In summary, the **RUPB-61** chemotype represents a promising lead for the development of a new class of orally active, long-acting, and resistance-resilient chemoprotective agents. Its dual-stage activity, favorable pharmacokinetics, and distinct mechanism of action position it well to fill a critical gap in the malaria elimination toolkit. With continued optimization and preclinical validation, PfPKG inhibitors may emerge as key components of integrated strategies for malaria prevention, treatment, and eventual eradication.

## Acknowledgments

This research was funded by W81XWH2010386 from the Department of Defense to P.B. and J.Y.R. This project has been funded in whole or in part with Federal funds from the National Institute of Allergy and Infectious Diseases, National Institutes of Health, Department of Health and Human Services, under Contract No. 75N93022C00036 (S.L., A.D.). This research used resources the NYX beamline 19-ID, supported by the New York Structural Biology Center, at the National Synchrotron Light Source II, a U.S. Department of Energy (DOE) Office of Science User Facility operated for the DOE Office of Science by Brookhaven National Laboratory under Contract No. DE-SC0012704. The NYX detector instrumentation was supported by grant S10OD030394 through the Office of the Director of the National Institutes of Health (K.P.B.). The funders had no role in study design, data collection and analysis, decision to publish, or preparation of the manuscript. The following reagents were obtained through BEI Resources, NIAID, NIH: *Plasmodium falciparum*, Strain K1, MRA-159, contributed by Dennis E. Kyle and strain 7G8, MRA-152, contributed by David Walliker. Strain TM90_C2B was a kind gift by Dr. Alison Roth, WRAIR. The pFastBac construct expressing P. vivax PKG was a kind gift from Dr. Halabelian and Dr. Aled Edwards at University of Toronto. We thank Alma Seitova for helpful discussion.

The material has been reviewed by the Walter Reed Army Institute of Research (WRAIR). There is no objection to its presentation and/or publication. The opinions or assertions contained herein are the private views of the authors and are not to be constructed as official or as reflecting the true views of the Department of the Army or the Department of Defense. Research was conducted under an IACUC-approved animal use protocol in an AAALAC International-accredited facility with a Public Health Service Animal Welfare Assurance and in compliance with the Animal Act and other federal statutes and regulations relating to laboratory animals. This research was supported in part by an appointment to the Department of Defense (DOD) Research Participation Program administered by the Oak Ridge Institute for Science and Education (ORISE) through an interagency agreement between the U.S. Department of Energy (DOE) and the DOD. ORISE is managed by ORAU under DOE contract number DE-SC0014664. All opinions expressed in this paper are the author’s and do not necessarily reflect the policies and views of DOD, DOE, or ORAU/ORISE.

## Methods

### Parasite Cultures and Growth Inhibition Assays

Asexual blood stage *P. falciparum* strains, including 3D7 and drug-resistant field isolates, were cultured using standard conditions in RPMI 1640 with 0.5% Albumax as described previously [25]. EC₅₀ values were determined using a SYBR Green I fluorescence assay after 72 h of compound exposure. EC_50_ values were derived by nonlinear regression in GraphPad Prism version 10.

### *P. falciparum* and *P. cynomolgi* Sporozoite Production and Isolation

Liver stage antimalarial activity was evaluated using sporozoites from *P. falciparum* (NF54 strain) and *P. cynomolgi bastianellii* (B strain). *P. falciparum*-infected *Anopheles stephensi* mosquitoes were reared at WRAIR, while *P. cynomolgi*-infected *Anopheles dirus* mosquitoes were obtained from the Armed Forces Research Institute of Medical Sciences (AFRIMS), Thailand. Sporozoites were isolated by dissecting salivary glands under sterile conditions, followed by gentle mechanical disruption. Freshly isolated sporozoites were immediately used for hepatocyte infection.

### Primary Hepatocyte Culture and Sporozoite Infection

Primary cryopreserved hepatocytes were selected based on host-parasite compatibility: primary simian hepatocytes (lot OQB) for *P. cynomolgi* and human hepatocytes (lot BGW) for *P. falciparum* (BioIVT Inc., Baltimore, MD, USA). Cells were thawed according to the manufacturer’s protocol and seeded into 384-well collagen-coated assay plates (Greiner Bio-One, Monroe, NC, USA) using InVitroGro CP plating medium (BioIVT) [36, 37]. Seeding densities were optimized for uniform monolayer formation, and hepatocytes were allowed to adhere for 24–48 hours before infection. Sporozoites were added at a 1:1.1 hepatocyte-to-sporozoite ratio within 48 hours post-seeding. Automated robotic systems were used for media changes and compound additions to ensure assay consistency and minimize variability associated with manual pipetting.

### Drug Treatment Regimens for P. falciparum and P. cynomolgi sporozoite assay

Test compounds (*Pf*PKG inhibitors) and assay drug controls (**KDU-691** and tafenoquine) were solubilized in DMSO or PBS and serially diluted into 8- or 12-point, 2- or 3-fold series, achieving final assay concentrations ranging from 20 µM to 0.019 µM. Compounds were introduced using a calibrated pin tool controlled by robotic arms (V&P Scientific), ensuring a 1:1000 dilution into assay wells. Media were replenished daily with fresh drug-containing HCM. Compounds were added 1-hour post-infection (p.i.) (day 0) and administered daily for 3 days. Cells were fixed on day 6 p.i. of *P. falciparum* culture and day 8 for *P. cynomolgi* infected culture, coinciding with liver stage schizont maturation.

### Fixation and Immunofluorescence Staining for *P. falciparum* and *P. cynomolgi* assays

Cells were fixed using 4% paraformaldehyde for 20 minutes at room temperature, followed by three PBS washes, and stored at 4°C until further processing. Parasite detection was performed using species-specific primary antibodies, with *P. cynomolgi* parasites labeled using a mouse monoclonal anti-*Plasmodium* GAPDH antibody (1.6 μg/mL, European Malaria Reagent Repository) and *P. falciparum* parasites detected using a rabbit polyclonal anti-PfHSP70 antibody (1:70 dilution, LS Bio, USA). Primary antibodies were diluted in PBS containing 1% bovine serum albumin (BSA) and 0.03% Triton X-100 and incubated overnight at 4°C. Following primary antibody incubation, cells were washed and incubated overnight at 4°C with Alexa Fluor Plus 555-conjugated goat anti-mouse/anti-rabbit IgG secondary antibodies (2 μg/mL, Invitrogen, USA). Nuclei were counterstained using Hoechst 33342 (5–10 μg/mL) for one hour at room temperature before imaging. Each assay run included a control plate containing uninfected hepatocytes stained under identical conditions.

### High-Content Imaging and Analysis

High-content imaging was performed using the Operetta CLS system (Revvity) and image analysis was performed with Harmony software version 4.9 (Revvity, Waltham, MA, USA). Images were acquired at 10x or 20x magnification using DAPI and TRITC fluorescence channels. Following the established protocol [38, 39], parasites were classified as hypnozoites or schizonts based on fluorescence intensity (mean and maximum), morphological features (cell roundness and size), and segmentation criteria. Hepatocyte nuclei were identified using Hoechst 33342 staining. Background signal was captured in uninfected, stained wells and subjected to the same parasite detection algorithm. Estimate erroneous parasite counts were then subtracted from infected well counts. Hepatocyte viability was determined by quantifying nuclear counts per well using Hoechst stain intensity thresholds. A total of 9 or 25 non-overlapping fields were captured per well, covering more than 95% of the total well surface area. An in-house built Python-based image analysis script identified parasite objects using predefined fluorescence intensity thresholds. Objects were validated based on size and shape criteria and confirmed via DAPI co-localization to eliminate false positives. Parasite size was quantified by calculating the average object area (µm²) of validated parasite structures per well. The classification pipeline was applied across all plates, accounting for debris and biological background estimated from uninfected stain controls. These values were subtracted from individual plate parasite counts to obtain normalized data. Dose-response inhibition and IC_50_ were calculated relative to in-plate negative controls (DMSO-treated wells). A four-parameter logistic regression model was fitted to the percentage survival or inhibition versus log concentration data using GraphPad Prism (Windows version 10.0.3, GraphPad Software Inc., USA) or a Python-based grid algorithm [40].

### Efficacy Studies

*In vivo* chemoprotective efficacy was evaluated using luciferase-expressing *P. berghei* ANKA sporozoites. Female Swiss-webster (Taconic) wild-type mice (6–8 weeks old) were administered **RUPB-61** (10 mg/kg in PBS) orally or intravenously prior to intravenous injection of 0.5 - 1 × 10^4^ sporozoites. Liver stage parasite burden was measured 42–48 h p.i. by bioluminescence imaging (IVIS Spectrum). Photon flux from the liver region was quantified using Living Image software. All animal procedures were approved by the Rutgers Institutional Animal Care and Use Committee (IACUC).

### Bioinformatic analysis of *P. falciparum* genomes

This study included sequences of *P. falciparum* generated by the MalariaGen consortium and available on the European Nucleotide Archive (https://www.ebi.ac.uk/ena). Whole-genome sequencing data were downloaded using the ena-file-downloader.jar tool to fetch paired reads in FASTQ format via FTP (File Transfer Protocol). Sequencing reads were trimmed for quality control using Sickle [41] with the default parameters (length threshold of 20 and Phred quality score threshold of 20). Trimmed files were then aligned to the *Pf3D7_01_v3* reference genome using bwa (v0.7.17-r1188). Samtools (v1.3.1) were used to generate BAM, sorted BAM, and index BAM files. Mutations were then identified from the aligned reads using Bcftools (v2.1) mpileup to generate genotype likelihoods at each genomic position with coverage and Bcftools call for variant calling. Variants were then annotated using SNPEff with Pf3D7_01_v3 as the reference genome. The resulting VCF files were compressed, indexed, and filtered by location (between 1490321 and 1494453 on chromosome 14). The VCF files were combined into one file for all samples. This final VCF file was downloaded and imported into R for further analysis. PfPKG sequence used is from positions 1490315 – 1494452 on chromosome 14.

### Pharmacokinetic Analysis

Mice were dosed with **RUPB-61** (10 mg/kg, P.O. or I.V.) and blood collected over 24 h. Plasma concentrations were measured by LC-MS/MS. Non-compartmental PK parameters were calculated using Phoenix WinNonlin. Human half-life predictions were derived by allometric scaling for half-life (t_1/2_): t_1/2_,human = t_1/2_,mouse × (W_human_/W_mouse_)^0.25^, where W = body weight. The exponent 0.25 is commonly used for time-based pharmacokinetic parameters. We assume an average mouse weight of 0.025 kg and an average human weight of 70 kg. With a mouse half-life of 9 -12 hours, we compute the human half-life range to be 67.5 – 90 h.

### Protein expression

*P. vivax* PKG was purified as previously described [42]. Briefly, Sf9 cells expressing the protein were flash frozen in liquid nitrogen. They were lysed hypotonically by thawing in 20 mM HEPES pH 7.0, 2 mM MgCl_2_, 0.5% Triton X-100 and EDTA-free protease inhibitor tablets (Roche). The whole cell lysate was flash frozen in liquid nitrogen. Thawed lysate was adjusted to 300 mM NaCl and 5% glycerol, and 1 mM TCEP, and insoluble material was removed by centrifugation at 10,000 RCF for 1 hour. The supernatant was treated with benzonase for 30 minutes, and debris pelleted at 230,000 rcf for 1 hour. Protein was purified by IMAC with an elution gradient of buffer A (20 mM HEPES pH 7.0, 300 mM NaCl, 1 mM TCEP, 5% glycerol) to buffer A with 500 mM Imidazole. Peak fractions were concentrated in a 10 kDa MWCO concentrator (Pall), then enriched by SEC on a 26/600 superdex 200 column (Cytiva) equilibrated in buffer A. Protein from peak fractions was concentrated to 10 mg/ml and flash frozen in liquid nitrogen.

### Crystallization

Purified PvPKG, in 20 mM HEPES pH 7.0, 300 mM NaCl, 5% (v/v) glycerol, 1 mM TCEP was diluted to 10 mg/mL for crystallization screening. Protein-ligand complexes were prepared by adding 2 mM **RUPB-60** and **RUPB-61**, from 100 mM DMSO stock solutions to aliquots of the protein and incubating on ice for 1 hour. All crystallization experiments were conducted using an NT8 drop-setting robot (Formulatrix Inc.) and UVXPO MRC (Molecular Dimensions) sitting drop vapor diffusion plates at 17 °C. 100 nL of protein and 100 nL crystallization solution were dispensed and equilibrated against 50 µL of the latter. Prismatic crystals were obtained overnight from the Morpheus [43] screen (Molecular Dimensions) condition B3 (20%(v/v) glycerol, 10% w/v PEG 4000, 100 mM Imidazole/MES, pH 6.5, 30 mM NaF, 30 mM NaBr and 30 mM NaI). Crystals were cryoprotected by layering the well solution onto the drops, harvesting directly and storing in liquid nitrogen. X-ray diffraction data were collected at the National Synchrotron Light Source II (NSLS-II) beamline 19-ID-I (NYX) using a Dectris Eiger2 9M XE pixel array detector. Data collection and refinement statistics are summarized in Supplementary Information.

### Structure solution and Refinement

Intensities were integrated using XDS [44, 45] via Autoproc [46] and the Laue class analysis and data scaling were performed with Aimless [47]. Structure solution was conducted by molecular replacement with Phaser [48] using a previously determined structure (PDB Intensities were integrated using XDS [44, 45] via Autoproc [46] and the Laue class analysis and data scaling were performed with Aimless [47]. Data from 6 crystals, obtained from the same crystallization condition, were scaled together to improve the data completeness and multiplicity due to the low symmetry *P*1 lattice. Structure solution was conducted by molecular replacement with Phaser [48] using a previously determined structure (PDB Intensities were integrated using XDS [44, 45] via Autoproc [46] and the Laue class analysis and data scaling were performed with Aimless [47]. Data from 6 crystals, obtained from the same crystallization condition, were scaled together to improve the data completeness and multiplicity due to the low symmetry *P*1 lattice. Structure solution was conducted by molecular replacement with Phaser [48] using a previously determined PfPKG structure (PDB 4RZ7) as the search model. Structure refinement, manual model building and validation were conducted with Phenix [49], Coot [50] and Molprobity [51] respectively.

### Accession Codes

The coordinates and structure factors for the Ba-Bfr structures have been deposited to the Worldwide Protein Databank (wwPDB.org) with the accession codes 9P75 (**RUPB-60**) and 9P74 (**RUPB-61**).

### Modeling

**RUPB-61** was docked into prepared PDB structures of ROCK1 using the 25-2 release of Schrödinger’s IFD-MD **[52]**. The representations of ROCK1 were PDB IDs 2ETR, 3TVJ and 2ETR with the c-term loop deleted. IFD-MD utilizes a combination of induced-fit docking and molecular dynamics to predict the pose of **RUPB-61** starting from the ROCK1 structures. Each predicted ligand-receptor complex produced by IFD-MD (15 poses in total for ROCK1) was then scored using Schrödinger’s FEP+ (Schrödinger Release 2025-2: FEP+, Schrödinger, LLC, New York, NY, 2025) using absolute binding free energy perturbation (AB-FEP) [53]. Due to finite simulation times, AB-FEP only accounts for a limited amount of the protein-reorganization free energy to transform the protein from the apo to holo state [53, 54]. To accurately predict absolute binding affinity, an offset was added to each AB-FEP prediction to correct for this missing free energy cost. We pair this offset with each of the starting PDB structures and apply it as a post-hoc correction to the AB-FEP predicted ΔGs. The final prediction is the most negative (favorable) AB-FEP ΔG after addition of the respective offset.

### Kinome Selectivity Profiling

Off-target kinase inhibition was assessed using the DiscoverX scanMAX (Eurofins) panel at 1 µM compound concentration. Binding affinities of CIT, ROCK1 and ROCK2 were evaluated using the KdElect assay (Eurofins).

### Synthetic Strategy

**Scheme 1.**
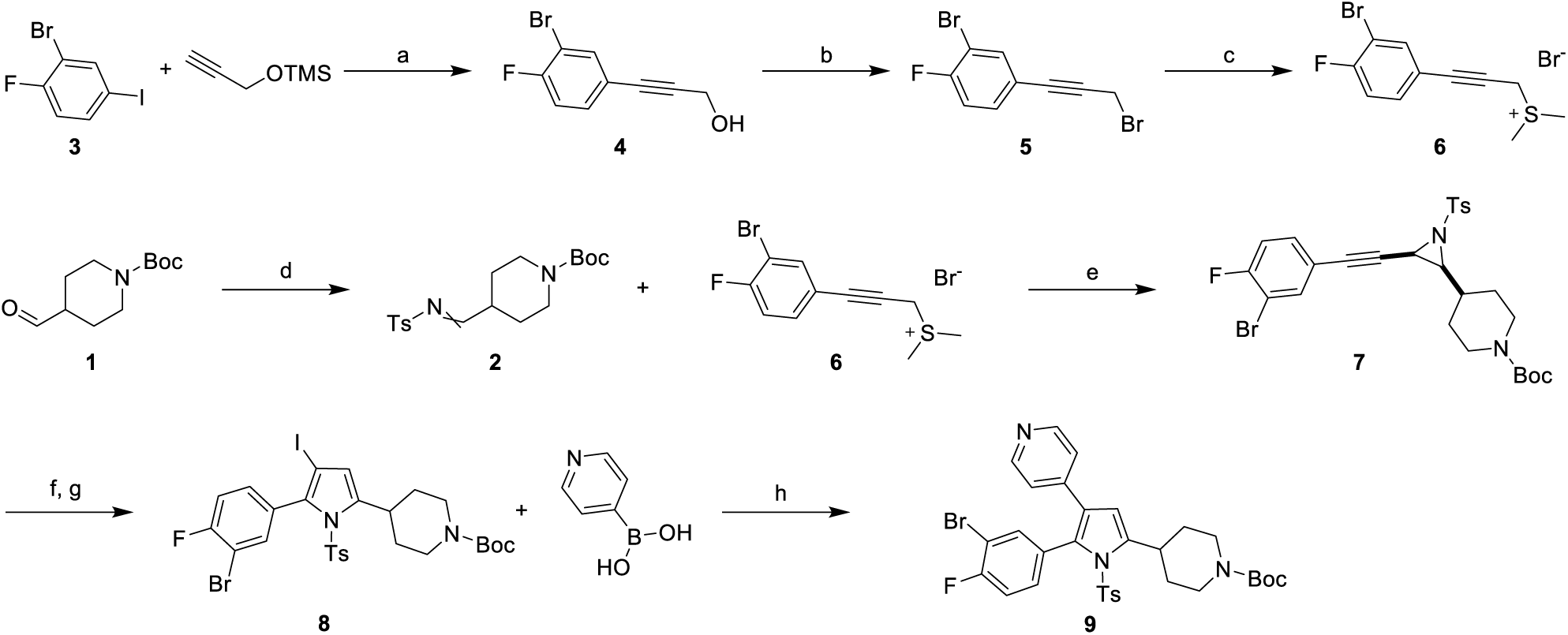
Synthesis of late-stage intermediate **9**. Reagents and conditions: a) CuI, Pd(PPh_3_)_2_Cl_2_, TEA, 20°C, 4 h; b) CBr_4_, PPh_3_, DCM, 20°C, 1 h; c) Me_2_S, acetone, 20°C, 12 h; d) TsNH_2_, PhSO_2_Na, HCOOH/H_2_O, 20°C, 12 h; e) Cs_2_CO_3_, DCM, 20°C, 1.5 h; f) PtCl_2_, I_2_, MeCN/H_2_O, 80°C, 30 min; g) Boc_2_O, TEA, DCM, 20°C, 1 h; h) Pd(dppf)Cl_2_·CH_2_Cl_2_, 1,4-dioxane, 2M Na_2_CO_3_ (aq), μW, 90°C, 20 min.

The synthesis of late-stage intermediates **9** and **12** begins with a Sonogashira coupling to afford alkyne **4**. Appel reaction with carbon tetrabromide to give **5** is followed by insertion of dimethyl sulfide, affording dimethylsulfonium salt **6**. Condensation of aldehyde **1** with p-toluene sulfonamide affords Schiff base **2**, that is then reacted with **6** to give cis-propargylic aziridine **7** under basic conditions. A platinum chloride catalyzed cycloisomerization in the presence of iodine affords the highly functionalized pyrrole **8**. Subsequently, Suzuki coupling with 4-pyridylboronic acid gives selective coupling at the iodide of **8** to give late-stage intermediate **9**. TBAF mediated tosyl deprotection of the pyrrole affords **10** that is then subjected to acidic deprotection conditions to give secondary amine **11** followed by reductive amination to arrive at late-stage intermediate **12**.

**Scheme 2.**
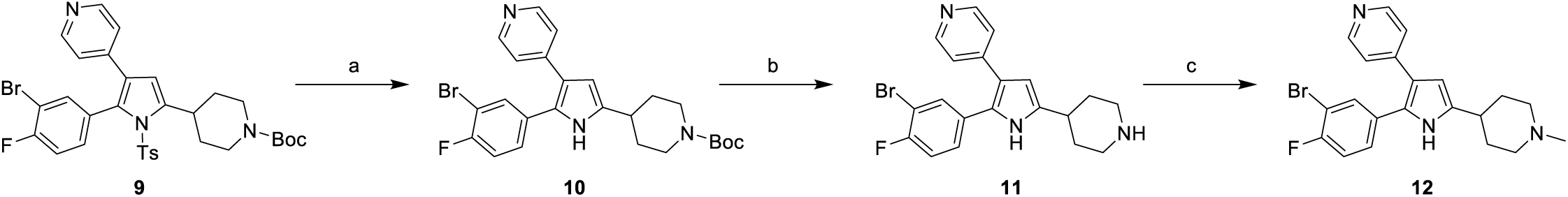
Synthesis of late-stage intermediate **12**. a) 1M TBAF/THF, THF, rt, 16 h; b) 6M HCl/IPA, DCM, MeOH; c) formalin, AcOH, MeOH then NaCNBH_3_, 1 h.

**Scheme 3.**
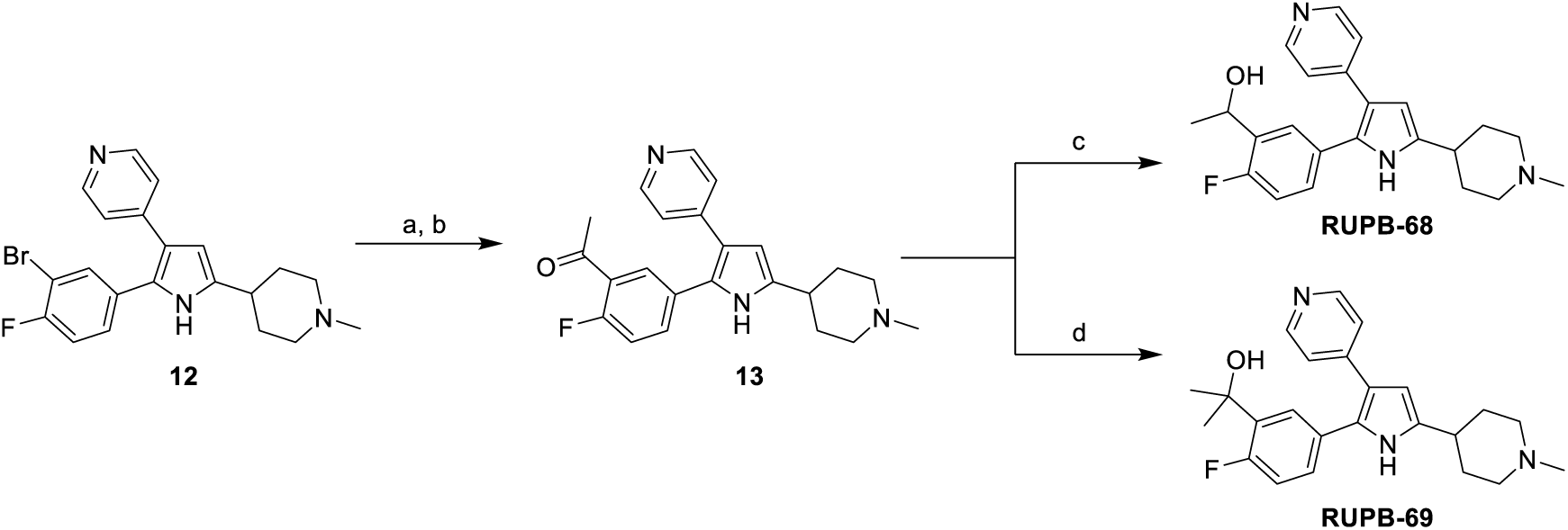
Synthesis of **RUPB-68** and **RUPB-69**. a) tributyl(1-ethoxyvinyl)stannane, PdCl_2_(PPh_3_)_2_, 1,4-dioxane, μW, 130°C, 30 min; b) 1M HCl(aq), THF, water, rt, 1 h; c) NaBH_4_, MeOH, rt, 5 min; d) 3M MeMgBr in Et_2_O, Et_2_O, rt, 2 h, then NH_4_Cl(aq), rt, 30 min.

Stille coupling of late-stage intermediate **12** and subsequent hydrolysis of the enone under acidic conditions gives **13**. Borohydride reduction of **13** gives racemic secondary alcohol **RUPB-68**. Methyl addition to the ketone of **13** with methyl Grignard reagent affords tertiary alcohol **RUPB-69**.

**Scheme 4.**
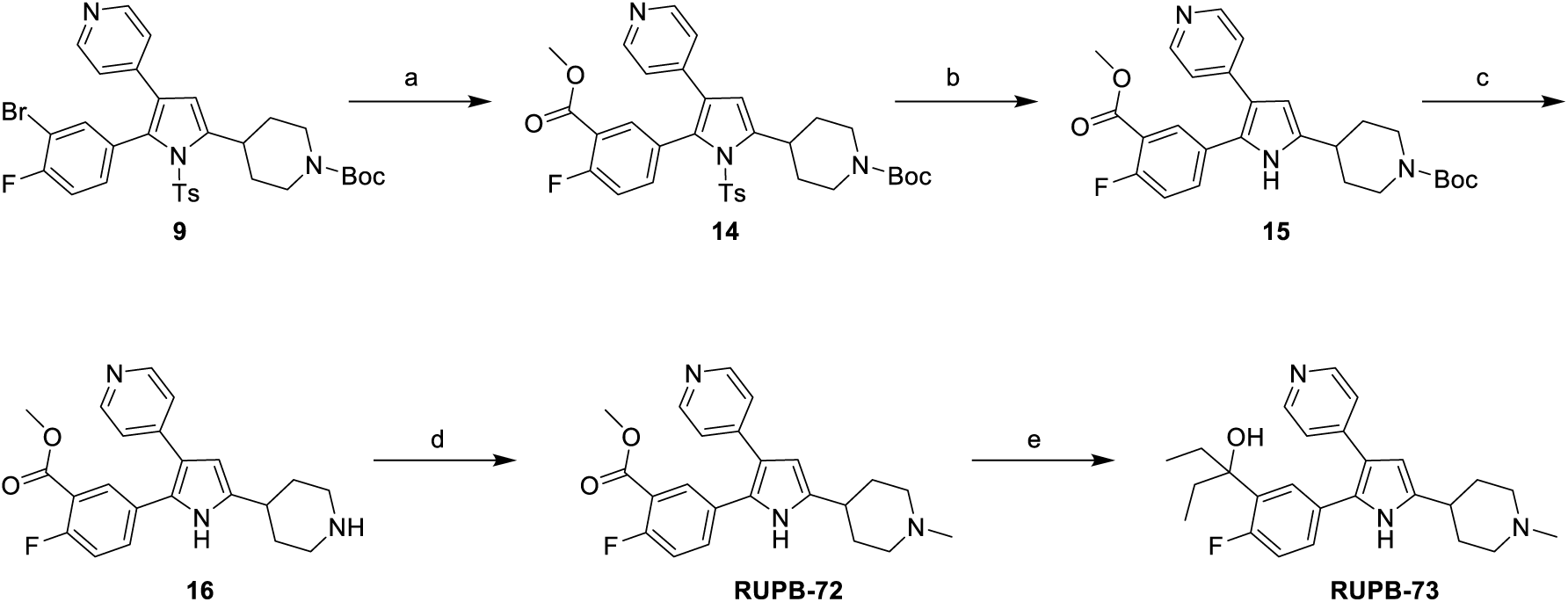
Synthesis of **RUPB-73**. a) NaOAc, Mo(CO)₆, Pd(dppf)Cl_2_·CH_2_Cl_2_, MeOH, μW, 140°C, 1 h; b) 1M TBAF/THF, THF, rt, 1 h; c) TFA, DCM, 30 min; d) formalin, AcOH, MeOH, then NaCNBH_3_, 1 h; e) 3M EtMgBr in Et_2_O, Ti(OiPr)_4_, THF, rt, 30 min.

Palladium catalyzed carbonylation into the bromide of intermediate **9** affords ester **14**. Using the previously described deprotection and methylation conditions, **14** is converted into **RUPB-72**. Diethyl insertion into the ester with ethylmagnesium bromide in the presence of titanium (IV) isopropoxide gives analog **RUPB-73**.

### Experimental Procedures for Chemical Synthesis and Characterization 3-(3-Bromo-4-fluorophenyl)prop-2-yn-1-ol (4)

To a mixture of 2-bromo-1-fluoro-4-iodobenzene **(3)** (40.0 g, 0.13 mol, 1.0 equiv.) and trimethyl(prop-2-yn-1-yloxy)silane (23.9 g, 0.186 mol, 1.43 equiv.) in THF (500 mL) was added TEA (20.1 g, 0.199 mol, 1.53 equiv.), copper(I) iodide (2.53 g, 0.013 mol, 0.10 equiv.) and Pd(PPh_3_)_2_Cl_2_ (4.66 g, 6.64 mmol, 0.05 equiv.). Then the reaction mixture was stirred under N_2_ at 20°C for 4 hours. TLC (Petroleum ether: Ethyl acetate = 10:1) showed that the starting material was consumed, and a new spot was detected. The mixture was diluted with saturated aq. NH_4_Cl (500 mL) and extracted with ethyl acetate (500 mL×3). The organic phase was concentrated under reduced pressure. The crude product was purified by silica gel chromatography (SiO_2_, Ethyl acetate/ Petroleum ether = 0-20%) to give 3-(3-bromo-4-fluorophenyl)prop-2-yn-1-ol **(4)** (23.0 g, 71.7% yield) as a colorless oil. ^1^H NMR (400 MHz, DMSO-*d*_6_) δ 7.78 (dd, *J* = 6.8, 2.0 Hz, 1H), 7.52 – 7.46 (m, 1H), 7.40 (t, *J* = 8.8 Hz, 1H), 5.37 (t, *J* = 6.0 Hz, 1H), 4.30 (d, *J* = 6.0 Hz, 2H).

### 2-Bromo-4-(3-bromoprop-1-yn-1-yl)-1-fluorobenzene (5)

To a solution of 3-(3-bromo-4-fluorophenyl)prop-2-yn-1-ol **(4)** (20.0 g, 87.3 mmol, 1.0 equiv.) in DCM (200 mL) was added CBr_4_ (34.7 g, 105 mmol, 1.2 equiv.). Then a solution of triphenylphosphine (27.5 g, 105 mmol, 1.2 equiv.) in DCM (200 mL) was added to the reaction mixture and stirred at 20°C for 1 hour. TLC (Petroleum ether: Ethyl acetate = 10:1) showed that the starting material was consumed, and a new spot was detected. The mixture was concentrated under reduced pressure. The crude product was diluted with n-hexanes (100 mL) and filtered. The filtrate was concentrated under reduced pressure to give 2-bromo-4-(3-bromoprop-1-yn-1-yl)-1-fluorobenzene **(5)** (24.0 g, 89.4% yield) as a colorless oil. ^1^H NMR (400 MHz, DMSO-*d*_6_) δ 7.84 (dd, *J* = 6.8, 2.0 Hz, 1H), 7.54 (ddd, *J* = 8.4, 4.8, 2.0 Hz, 1H), 7.42 (t, *J* = 8.8 Hz, 1H), 4.51 (s, 2H).

### (3-(3-Bromo-4-fluorophenyl)prop-2-yn-1-yl)dimethylsulfonium bromide (6)

To a solution of 2-bromo-4-(3-bromoprop-1-yn-1-yl)-1-fluorobenzene **(5)** (24.0 g, 0.12 mol, 1 equiv.) in acetone (240 mL) was added Me_2_S (23.0 g, 0.37 mol, 3.09 equiv.). The reaction mixture was stirred at 20°C for 12 hours. The mixture was filtered, and the solid was collected and dried under vacuum to give (3-(3-bromo-4-fluorophenyl)prop-2-yn-1-yl)dimethylsulfonium bromide **(6)** (30.0 g, 90.0% yield) as a white solid. ^1^H NMR (400 MHz, DMSO-*d*_6_) δ 8.08 (dd, *J* = 6.8, 2.0 Hz, 1H), 7.68 (ddd, *J* = 8.4, 4.8, 2.0 Hz, 1H), 7.47 (t, *J* = 8.8 Hz, 1H), 4.70 (d, *J* = 13.6 Hz, 2H), 3.00 – 2.89 (m, 6H).

### *tert*-Butyl 4-((tosylimino)methyl)piperidine-1-carboxylate (2)

To a mixture of *tert*-butyl 4-formylpiperidine-1-carboxylate **(1)** (55.0 g, 0.257 mol, 1 equiv.) and sodium p-toluenesulfinate (50.3 g, 0.28 mol, 1.09 equiv.) in FA/H_2_O (800 mL, 1: 1) was added 4-methylbenzenesulfonamide (44.0 g, 0.257 mol, 1 equiv.) The reaction mixture was stirred at 20°C for 12 hours. TLC (SiO_2_, Petroleum ether: Ethyl acetate = 1:1) showed that the starting material was consumed, and a new spot was detected. The mixture was filtered, and the filter cake was washed with water (500 mL×2) and hexanes (500 mL×2). The solid was dried under vacuum to give *tert*-butyl 4-((tosylimino)methyl)piperidine-1-carboxylate **(2)** (60.0 g, 0.154 mol, 60.2% yield, 100% purity by LCMS) as a white solid. ((MH-100)+) 267.10.

### *tert*-Butyl 4-((2*R*,3*S*)-3-((3-bromo-4-fluorophenyl)ethynyl)-1-tosylaziridin-2-yl)piperidine-1-carboxylate (7)

To a solution of *tert*-butyl 4-((tosylimino)methyl)piperidine-1-carboxylate **(2)** (15.0 g, 40.7 mmol, 1 equiv.) in DCM (300 mL) was added (3-(3-bromo-4-fluorophenyl)prop-2-yn-1-yl)dimethylsulfonium bromide **(6)** (14.4 g, 40.7 mmol, 1 equiv.) and Cs_2_CO_3_ (13.3 g, 40.7 mmol, 1 equiv.). Then the reaction mixture was stirred at 20°C for 1.5 hours. LCMS showed that the starting material was consumed, and the desired product was detected as a major peak. The mixture was diluted with saturated aq. NH_4_Cl (200 mL) and extracted with DCM (200 mL×2). The organic phase was concentrated under reduced pressure. The crude product was purified by silica gel chromatography (SiO_2_, Ethyl acetate/ Petroleum ether = 0-20%) to give *tert*-butyl 4-((2*R*,3*S*)-3-((3-bromo-4-fluorophenyl)ethynyl)-1-tosylaziridin-2-yl)piperidine-1-carboxylate **(7)** (20.0 g, 80.5% yield, 100% purity by LCMS) as a yellow solid. (MNa+) 599.05.

### *tert*-Butyl 4-(5-(3-bromo-4-fluorophenyl)-4-iodo-1-tosyl-1H-pyrrol-2-yl)piperidine-1-carboxylate (8)

To a solution of *tert*-butyl 4-((2*R*,3*S*)-3-((3-bromo-4-fluorophenyl)ethynyl)-1-tosylaziridin-2-yl)piperidine-1-carboxylate **(7)** (20.0 g, 34.5 mmol, 1.0 equiv.) in MeCN/H_2_O (880 mL, v/v = 10: 1) was added I_2_ (17.5 g, 69.0 mmol, 1.5 equiv.) and PtCl_2_ (1.38 g, 5.17 mmol, 0.15 equiv.). The reaction mixture was heated to 80°C for 30 minutes. The mixture was diluted with 10% aq. Na_2_SO_3_ (800 mL) and extracted with ethyl acetate (800 mL×2). The organic phase was concentrated under reduced pressure. The crude product was diluted with DCM (200 mL) followed by addition of Boc_2_O (8.28 g, 37.9 mmol) and TEA (6.97 g, 69.0 mmol) to the solution. Then the mixture was stirred at 20°C for 1 hour. LCMS showed that 30% starting material remaining with 40% desired product. The mixture was concentrated under reduced pressure. The crude product was purified by silica gel chromatography (SiO_2_, Ethyl acetate/ Petroleum ether = 0-15%) to give *tert*-butyl 4-(5-(3-bromo-4-fluorophenyl)-4-iodo-1-tosyl-1H-pyrrol-2-yl)piperidine-1-carboxylate **(8)** (4.00 g, 15.6% yield, 98.6% purity by HPLC) as a white solid. (MNa)+ 725.05. ^1^H NMR (400 MHz, DMSO-*d*_6_) δ 7.43 – 7.32 (m, 5H), 7.26 (dd, *J* = 6.4, 1.6 Hz, 1H), 7.22 (ddd, *J* = 8.4, 4.8, 2.0 Hz, 1H), 6.46 (s, 1H), 4.13 – 3.94 (m, 2H), 3.33 – 3.26 (m, 1H), 2.93 – 2.69 (m, 2H), 2.39 (s, 3H), 1.87 – 1.75 (m, 2H), 1.46 – 1.37 (m, 11H). ^19^F NMR (400 MHz, DMSO-*d*_6_) δ -108.03 (s, 1F).

### *tert*-Butyl 4-(5-(3-bromo-4-fluorophenyl)-4-(pyridin-4-yl)-1-tosyl-1H-pyrrol-2-yl)piperidine-1-carboxylate (9)

A mixture of *tert*-butyl 4-(5-(3-bromo-4-fluorophenyl)-4-iodo-1-tosyl-1H-pyrrol-2-yl)piperidine-1-carboxylate **(8)** (700 mg, 1.0 mmol, 1 equiv.), pyridin-4-ylboronic acid (148 mg, 1.2 mmol, 1.2 equiv.) and Pd(dppf)Cl_2_·CH_2_Cl_2_ (82 mg, 0.10 mmol, 0.10 equiv.) was taken up in 1,4-dioxane (10 mL) and 2M Na_2_CO_3_ (aq) (5 mL) was heated in a microwave reactor for 20 minutes at 90°C. The reaction was determined to be complete by LCMS with a 73:9 ratio of mono-addition:double-addition products by LCMS at 254 nm. The crude reaction mixture was partitioned in EtOAc/water. The aqueous was extracted 3 x EtOAc. The combined organic was washed 1 × brine, dried over sodium sulfate, filtered and concentrated. The concentrate was purified by silica gel chromatography (0% → 5% MeOH/DCM) and product fractions repurified by silica gel chromatography (0% → 50% → 100% EtOAc/Hex). Product fractions were concentrated under reduced pressure and dried under vacuum to afford *tert*-butyl 4-(5-(3-bromo-4-fluorophenyl)-4-(pyridin-4-yl)-1-tosyl-1H-pyrrol-2-yl)piperidine-1-carboxylate **(9)** as a white solid (498 mg, 95% purity by LCMS). (MH+) 654.30, 656.30.

### *tert*-Butyl 4-(5-(3-bromo-4-fluorophenyl)-4-(pyridin-4-yl)-1H-pyrrol-2-yl)piperidine-1-carboxylate (10)

To a solution of *tert*-butyl 4-(5-(3-bromo-4-fluorophenyl)-4-(pyridin-4-yl)-1-tosyl-1H-pyrrol-2-yl)piperidine-1-carboxylate **(9)** (389 mg, 0.59 mmol, 1 equiv.) in THF (20 mL) was added TBAF (1M in THF) (2.95 mL, 2.95 mmol, 5 equiv.). After stirring for 16 hours at room temperature, the reaction was determined complete by LCMS and was concentrated under reduced pressure. The residue was partitioned in EtOAc/water. The aqueous was separated and the organic was washed 1 × brine, dried over sodium sulfate, filtered and concentrated. The residue was purified by silica gel chromatography (0% → 2% MeOH/DCM). The combined product containing fractions were concentrated under reduced pressure. The solid was taken up in EtOAc, sonicated and filtered. The solid was washed with minimal EtOAc and dried under vacuum to yield *tert*-butyl 4-(5-(3-bromo-4-fluorophenyl)-4-(pyridin-4-yl)-1H-pyrrol-2-yl)piperidine-1-carboxylate **(10)** (258.8 mg, 0.52 mmol, 98% purity by LCMS) as a tan solid. (MH+) 500.20, 502.20.

### 4-(2-(3-Bromo-4-fluorophenyl)-5-(piperidin-4-yl)-1H-pyrrol-3-yl)pyridine (HCl salt) (11)

To a solution of tert-butyl 4-[5-(3-bromo-4-fluorophenyl)-4-(pyridin-4-yl)-1H-pyrrol-2-yl]piperidine-1-carboxylate **(10)** (258.8 mg, 0.52 mmol, 1 equiv.) in DCM (5 mL) and MeOH (5 mL) was added 6M HCl in IPA (1 mL) and stirred for 16 hours at rt. The reaction was determined to be complete by LCMS and concentrated under reduced pressure to give 4-(2-(3-bromo-4-fluorophenyl)-5-(piperidin-4-yl)-1H-pyrrol-3-yl)pyridine (HCl salt) **(11)** in quantitative yield (97% purity by LCMS at 254 nm). (MH+) 400.15, 402.20.

### 4-(2-(3-Bromo-4-fluorophenyl)-5-(1-methylpiperidin-4-yl)-1H-pyrrol-3-yl)pyridine (12)

To a solution of 4-[2-(3-bromo-4-fluorophenyl)-5-(piperidin-4-yl)-1H-pyrrol-3-yl]pyridine **(11)** (158 mg, 0.33 mmol, 1.0 equiv.) in MeOH (10 ml) was added AcOH (400 µL) and formalin (800 µL). After stirring for 1 hour at room temperature, an excess of sodium cyanoborohydride was added. Starting material and product coelute by LCMS. The reaction was determined to be complete when scan by scan analysis of the product peak showed only mass for methylated product with no visible signal for starting material mass present. After stirring for 30 minutes post addition of sodium cyanoborohydride, the reaction was concentrated under reduced pressure to remove the majority of the methanol. The residue was partitioned in EtOAc/water. The product was extracted into the acidic aqueous with 2 x water extractions of the EtOAc layer. The combined acidic aqueous was neutralized with NaHCO_3_ (sat, aq) and extracted 3 x DCM. The combined organic was dried over sodium sulfate, filtered and concentrated to afford 4-(2-(3-bromo-4-fluorophenyl)-5-(1-methylpiperidin-4-yl)-1H-pyrrol-3-yl)pyridine **(12)** as a yellow solid (134 mg, 99% purity by LCMS at 254 nm). (MH+) 414.10, 415.85.

### 1-(2-Fluoro-5-(5-(1-methylpiperidin-4-yl)-3-(pyridin-4-yl)-1H-pyrrol-2-yl)phenyl)ethan-1-one (13)

A mixture of 4-(2-(3-bromo-4-fluorophenyl)-5-(1-methylpiperidin-4-yl)-1H-pyrrol-3-yl)pyridine **(12)** (34.2 mg, 0.083 mmol, 1 equiv.), tributyl(1-ethoxyvinyl)stannane (74.5 mg, 0.21 mmol, 2.5 equiv.) and PdCl_2_(PPh_3_)_2_ (5.8 mg, 0.0083 mmol, 0.10 equiv.) was taken up in 1,4-dioxane (4 mL) and heated in a microwave reactor for 30 minutes at 130°C. The reaction was determined complete, as a mixture of the title compound and the enol intermediate. The mixture was diluted in EtOAc and filtered through celite to remove insoluble impurities. The filtrate was concentrated and taken up in a mixture of THF (4 mL) and water (4 mL). To this mixture was added 1M HCl(aq) (0.5 mL). After stirring for 1 hour at room temperature the reaction was determined complete by LCMS, with full conversion to the desired ketone product. The reaction was partitioned in DCM/water. The product partitioned into the acidic aqueous and the aqueous was washed 2 x DCM. The aqueous was basified with 1M NaOH(aq) and extracted 4 x DCM. The combined organic was dried over sodium sulfate, filtered and concentrated. The product was 98.5% purity after extraction but was purified by amino-d silica gel chromatography (0% → 2% MeOH/DCM) to remove any remaining impurities. The product fractions were combined and concentrated to afford 1-(2-fluoro-5-(5-(1-methylpiperidin-4-yl)-3-(pyridin-4-yl)-1H-pyrrol-2-yl)phenyl)ethan-1-one **(13)** (28.7 mg, 98% purity by LCMS at 254 nm) as a tan solid. (MH+) 378.30.

### 2-(2-Fluoro-5-(5-(1-methylpiperidin-4-yl)-3-(pyridin-4-yl)-1H-pyrrol-2-yl)phenyl)propan-2-ol (TFA salt) (RUPB-69)

To 1-(2-fluoro-5-(5-(1-methylpiperidin-4-yl)-3-(pyridin-4-yl)-1H-pyrrol-2-yl)phenyl)ethan-1-one **(13)** (15 mg, 0.040 mmol, 1 equiv.) in an oven-dried flask was added Et_2_O (3.5 mL). To this suspension was added MeMgBr (3M in Et_2_O) (60 µL, 0.20 mmol, 5 equiv.). After stirring for 1 hour at room temperature the reaction was roughly 45% complete by LCMS. An additional batch of MeMgBr (3M in Et_2_O) (60 µL, 0.20 mmol, 5 equiv.) was added. The reaction was roughly 67% complete by LCMS but was beginning to generate additional impurities. The reaction was quenched with NH_4_Cl (sat, aq) and was stirred for 30 min at rt. The mixture was partitioned in EtOAc/water and the aqueous was adjusted to pH = 2 with 1M HCl. The organic was extracted 2 x water. The combined aqueous was basified with 1M HCl and extracted 4 x DCM. The combined organic from the basic extraction was dried over sodium sulfate, filtered and concentrated. The concentrate was purified by prep HPLC (0.1% TFA(aq)/0.1% TFA(MeCN). The combined product fractions were concentrated and lyophilized to give 2-(2-fluoro-5-(5-(1-methylpiperidin-4-yl)-3-(pyridin-4-yl)-1H-pyrrol-2-yl)phenyl)propan-2-ol (TFA salt) **(RUPB-69)** (10.6 mg, 99% purity by LCMS) as a yellow/orange solid. (MH+) 394.30. ^1^H NMR (500 MHz, DMSO-*d*_6_) δ 11.89 – 11.72 (m, 1H), 10.37 – 10.17 (m, 1H), 8.63 – 8.51 (m, 2H), 7.73 – 7.65 (m, 3H), 7.34 (ddd, *J* = 8.3, 4.6, 2.4 Hz, 1H), 7.29 – 7.21 (m, 1H), 6.75 – 6.51 (m, 1H), 3.51 (d, *J* = 12.0 Hz, 2H), 3.17 – 3.03 (m, 2H), 2.91 – 2.73 (m, 4H), 2.28 – 2.17 (m, 2H), 1.87 (qd, *J* = 13.3, 3.7 Hz, 2H), 1.54 – 1.47 (m, 6H).

### 1-(2-Fluoro-5-(5-(1-methylpiperidin-4-yl)-3-(pyridin-4-yl)-1H-pyrrol-2-yl)phenyl)ethan-1-ol (TFA salt) (RUPB-68)

To a solution of 1-(2-fluoro-5-(5-(1-methylpiperidin-4-yl)-3-(pyridin-4-yl)-1H-pyrrol-2-yl)phenyl)ethan-1-one **(13)** (8 mg, 0.021 mmol, 1 equiv.) in MeOH (2.5 mL) was added NaBH_4_ (8 mg, 0.21 mmol, 10 equiv.). After stirring for 5 minutes at room temperature, the reaction was determined complete by LCMS. The reaction was concentrated under reduced pressure and the residue was purified by prep HPLC (0.1% TFA(aq)/0.1% TFA(MeCN). The combined product fractions were concentrated under reduced pressure and lyophilized to give 1-(2-fluoro-5-(5-(1-methylpiperidin-4-yl)-3-(pyridin-4-yl)-1H-pyrrol-2-yl)phenyl)ethan-1-ol (TFA salt) **(RUPB-68)** (8.2 mg, 99% purity by LCMS) as a yellow/orange solid. (MH+) 380.40. ^1^H NMR (500 MHz, DMSO-*d*_6_) δ 11.84 – 11.70 (m, 1H), 10.51 – 10.22 (m, 1H), 8.67 – 8.47 (m, 2H), 7.67 (d, *J* = 6.1 Hz, 2H), 7.56 (dd, *J* = 7.1, 2.4 Hz, 1H), 7.35 (ddd, *J* = 7.7, 4.9, 2.4 Hz, 1H), 7.26 (dt, *J* = 10.1, 7.0 Hz, 1H), 6.71 – 6.49 (m, 1H), 5.01 (q, *J* = 6.4 Hz, 1H), 3.50 (d, *J* = 12.2 Hz, 2H), 3.08 (q, *J* = 11.1 Hz, 2H), 2.90 – 2.74 (m, 4H), 2.24 (d, *J* = 13.2 Hz, 2H), 1.90 (tt, *J* = 14.6, 7.8 Hz, 2H), 1.34 (d, *J* = 6.3 Hz, 3H).

### *tert*-Butyl 4-(5-(4-fluoro-3-(methoxycarbonyl)phenyl)-4-(pyridin-4-yl)-1-tosyl-1H-pyrrol-2-yl)piperidine-1-carboxylate (14)

A mixture of tert-butyl 4-(5-(3-bromo-4-fluorophenyl)-4-(pyridin-4-yl)-1-tosyl-1H-pyrrol-2-yl)piperidine-1-carboxylate **(9)** (156 mg, 0.24 mmol, 1 equiv.), sodium acetate (196 mg, 2.4 mmol, 10 equiv.), molybdenum hexacarbonyl (253 mg, 0.96 mmol, 4 equiv.) and Pd(dppf)Cl_2_·CH_2_Cl_2_ (39 mg, 0.048 mmol, 0.20 equiv.) was taken up in MeOH (6 mL) and heated in a microwave reactor for 1 hour at 130°C. The reaction was incomplete by LCMS and heated in the microwave reactor an additional 40 minutes at 140°C. The reaction was partitioned in DCM/water and filtered over celite to remove insoluble solids. The product was rinsed through the filter with DCM/water. The layers were separated and the aqueous was extracted 2 x DCM. The combined organic was dried over sodium sulfate, filtered and concentrated to afford a mixture of *tert*-butyl 4-(5-(4-fluoro-3-(methoxycarbonyl)phenyl)-4-(pyridin-4-yl)-1H-pyrrol-2-yl)piperidine-1-carboxylate **(15)** (24% area of LCMS at 254 nm, (MH+) 480.30) and *tert*-butyl 4-(5-(4-fluoro-3-(methoxycarbonyl)phenyl)-4-(pyridin-4-yl)-1-tosyl-1H-pyrrol-2-yl)piperidine-1-carboxylate **(14)** (42% area of LCMS at 254 nm, (MH+) 634.35) that was carried forward to the next reaction without further purification.

### *tert*-Butyl 4-(5-(4-fluoro-3-(methoxycarbonyl)phenyl)-4-(pyridin-4-yl)-1H-pyrrol-2-yl)piperidine-1-carboxylate (15)

To the crude mixture of tert-butyl 4-(5-(4-fluoro-3-(methoxycarbonyl)phenyl)-4-(pyridin-4-yl)-1-tosyl-1H-pyrrol-2-yl)piperidine-1-carboxylate **(14)** and tert-butyl 4-(5-(4-fluoro-3-(methoxycarbonyl)phenyl)-4-(pyridin-4-yl)-1H-pyrrol-2-yl)piperidine-1-carboxylate **(15)** from the previous reaction in THF (10 mL) was added TBAF (1M in THF, 1 mL). The reaction was determined complete by LCMS after stirring for 1 hour at room temperature and was concentrated under reduced pressure to reduce the volume of THF. The concentrate was partitioned in EtOAc/water and the aqueous was extracted 3 x EtOAc. The combined organic was dried over sodium sulfate, filtered and concentrated. The concentrate was purified by silica gel chromatography (0% → 3% MeOH/DCM) and product fractions were combined, concentrated and dried under vacuum to afford *tert*-butyl 4-(5-(4-fluoro-3-(methoxycarbonyl)phenyl)-4-(pyridin-4-yl)-1H-pyrrol-2-yl)piperidine-1-carboxylate **(15)** as an orange solid (68.3 mg, 91% purity by LCMS at 254 nm). (MH+) 480.30.

### Methyl 2-fluoro-5-(5-(piperidin-4-yl)-3-(pyridin-4-yl)-1H-pyrrol-2-yl)benzoate (TFA salt) (16)

To a solution of *tert*-butyl 4-(5-(4-fluoro-3-(methoxycarbonyl)phenyl)-4-(pyridin-4-yl)-1H-pyrrol-2-yl)piperidine-1-carboxylate **(15)** (68.3 mg, 0.14 mmol) in DCM (2 mL) was added TFA (0.5 mL). The reaction was determined complete by LCMS after stirring 30 minutes at room temperature. The crude reaction was concentrated to afford methyl 2-fluoro-5-(5-(piperidin-4-yl)-3-(pyridin-4-yl)-1H-pyrrol-2-yl)benzoate (TFA salt) **(16)** (quantitative yield, yellow/orange solid) that was used in the next reaction without further purification. (MH+) 380.15.

### Methyl 2-fluoro-5-(5-(1-methylpiperidin-4-yl)-3-(pyridin-4-yl)-1H-pyrrol-2-yl)benzoate (RUPB-72)

To a solution of methyl 2-fluoro-5-(5-(piperidin-4-yl)-3-(pyridin-4-yl)-1H-pyrrol-2-yl)benzoate 2 x TFA (**16**) (85 mg, 0.14 mmol) in MeOH (4 mL) was added AcOH (200 µL) followed by formalin (400 µL). After stirring for 1 hour at room temperature, the reaction was quenched with the addition of excess sodium cyanoborohydride. The reaction was stirred for 30 minutes at room temperature and determined complete by LCMS. The crude reaction was concentrated under reduced pressure and the concentrate was partitioned in DCM and NaHCO_3_ (sat, aq). The aqueous was extracted 3 x DCM and the combined organic was dried over sodium sulfate, filtered, and concentrated. The product was purified by prep HPLC (0.1% FA(aq), MeCN(unmodified)). Product fractions were combined and concentrated to afford the title compound (64.6 mg yellow solid, formate salt, 80% purity by LCMS, (MH+) 394.15). The product was converted to the free base for solubility reasons related to the next reaction. The product was partitioned in DCM and NaHCO_3_ (sat, aq) and extracted 3 x DCM. The combined organic was dried over sodium sulfate, filtered and concentrated to give methyl 2-fluoro-5-(5-(1-methylpiperidin-4-yl)-3-(pyridin-4-yl)-1H-pyrrol-2-yl)benzoate **(RUPB-72)** as the free base (28.3 mg white solid, 80% purity by LCMS at 254 nm). (MH+) 394.15.

### 3-(2-Fluoro-5-(5-(1-methylpiperidin-4-yl)-3-(pyridin-4-yl)-1H-pyrrol-2-yl)phenyl)pentan-3-ol (TFA salt) (RUPB-73)

To a solution of methyl 2-fluoro-5-(5-(1-methylpiperidin-4-yl)-3-(pyridin-4-yl)-1H-pyrrol-2-yl)benzoate **(RUPB-72)** (20 mg, 0.051 mmol, 1 equiv.) in THF (4 mL) was added titanium(IV) isopropoxide (73 µL, 0.25 mmol, 5 equiv.) followed by ethylmagnesium bromide (3M in Et_2_O, 170 µL, 0.51 mmol, 10 equiv.). The reaction was stirred for 30 minutes at room temperature and quenched with water. The crude mixture was partitioned in EtOAc/water and filtered to remove solids. The product was rinsed through the filter with additional EtOAc/water. The organic was separated and the aqueous was extracted 2 x EtOAc. The combined organic was dried over sodium sulfate, filtered and concentrated. The concentrate was purified by prep HPLC (0.1% FA(aq)/MeCN(unmodified)). The crude reaction contained a mixture of unreacted starting material and the product of diethyl addition into the ester. Fractions containing pure products were combined separately and concentrated. Each of the products were then separately repurified by prep HPLC (0.1% TFA(aq)/0.1%TFA(MeCN). Clean fractions for each product were separately combined and concentrated to give the title compounds as TFA salts after concentration. Methyl 2-fluoro-5-(5-(1-methylpiperidin-4-yl)-3-(pyridin-4-yl)-1H-pyrrol-2-yl)benzoate (TFA salt) **(RUPB-72)** (5.8 mg sticky yellow/orange solid, 96.4% purity by HPLC at 254 nm. (MH+) 394.25. 3-(2-Fluoro-5-(5-(1-methylpiperidin-4-yl)-3-(pyridin-4-yl)-1H-pyrrol-2-yl)phenyl)pentan-3-ol (TFA salt) **(RUPB-73)** (9.6 mg sticky yellow/orange solid, 100% purity by HPLC at 254 nm). (MH+) 422.25. ^1^H NMR (500 MHz, DMSO-*d*_6_) δ 11.87 – 11.69 (m, 1H), 10.41 – 10.26 (m, 1H), 8.54 – 8.47 (m, 2H), 7.61 (d, *J* = 6.4 Hz, 2H), 7.54 (dd, *J* = 7.7, 2.5 Hz, 1H), 7.38 (ddd, *J* = 8.2, 4.6, 2.4 Hz, 1H), 7.27 (dt, *J* = 11.7, 7.6 Hz, 1H), 6.73 – 6.49 (m, 1H), 4.69 (s, 1H), 3.50 (d, *J* = 12.1 Hz, 2H), 3.08 (q, *J* = 11.6 Hz, 2H), 2.89 – 2.75 (m, 4H), 2.24 (d, *J* = 13.4 Hz, 2H), 1.92 (dddd, *J* = 22.5, 15.9, 10.9, 5.5 Hz, 4H), 1.70 (dq, *J* = 14.5, 7.3 Hz, 2H), 0.66 (t, *J* = 7.3 Hz, 6H).

**RUPB-58, -59, -60 and -61** were synthesized as described previously [25].

**RUPB-58** 4-{5-[(3*R*,4*R*)-1,3-Dimethylpiperidin-4-yl]-2-(4-fluorophenyl)-1H-pyrrol-3-yl}pyridine Dihydrochloride

**RUPB-59** 4-{5-[(3*R*,4*S*)-1,3-Dimethylpiperidin-4-yl]-2-(4-fluorophenyl)-1H-pyrrol-3-yl}pyridine Dihydrochloride

**RUPB-60** 4-{5-[(3*S*,4*R*)-1,3-Dimethylpiperidin-4-yl]-2-(4-fluorophenyl)-1H-pyrrol-3-yl}pyridine Dihydrochloride

**RUPB-61** 4-{5-[(3*S*,4*S*)-1,3-Dimethylpiperidin-4-yl]-2-(4-fluorophenyl)-1H-pyrrol-3-yl}pyridine Dihydrochloride

## Supplementary Figure Legends

**Supplementary Figure 1: Activity and cytotoxicity of PfPKG inhibitors in primary hepatocyte models of *P. falciparum and P. cynomolgi* sporozoite infection. (A)** Dose-response curve of **TSP** against *P. cynomolgi* hepatic hypnozoites formed in simian hepatocytes. Tafenoquine was used as a positive control. Data shown are normalized to number of infected cells in vehicle control. Data shown are from a single trial performed in technical duplicates. **(B)** Dose-response curves from two independent trials of **RUPB-61** against *P. falciparum* sporozoites in human hepatocytes. Data shown are normalized to the number of infected cells in vehicle control. **KDU-691** was used as a positive control. (**C)** Effect of **TSP** on the survival of primary simian hepatocytes. Data shown are from a single biological replicate performed in technical duplicates. **(D)** Effect of **RUPB-61** on the survival of primary human hepatocytes. Data shown are mean of two independent trials performed in technical duplicates. **(E)** Dose-response curves from three independent trials of **RUPB-61** against *P. cynomolgi* sporozoites in primary simian hepatocytes. Data shown are normalized to the number of infected cells in vehicle control. **KDU-691** was used as a positive control. **(F)** Effect of **RUPB-61** on the survival of primary simian hepatocytes. Data shown are mean of three independent trials performed in technical duplicates or triplicates.

**Supplementary Figure 2: Dose-response curves of RUPB-58 to -61 against multidrug resistant *P. falciparum* parasites.** Growth inhibition of 3D7, Dd2-B2, TM90_C2B, K1, 7G8 over 72h in asexual growth assays was measured using SybrGreen incorporation.

**Supplementary Figure 3: Liver parasitemia after administration of PfPKG inhibitors.** Bioluminescent imaging of mice to quantify liver parasitemia in mice infected with PbLuc-GFP sporozoites. **(A)** Mice administered three intravenous doses of vehicle or compound. Images were obtained at 44 hpi and 144 hpi. **(B)** Mice administered three oral doses of vehicle or compound. Images were obtained at 44 hpi and 144 hpi.

**Supplementary Figure 4: TREE*spot* Interaction map for RUPB-61.** Interaction of **RUPB-61** at 1 µM with 400 unique human kinases was determined using a panel of *in vitro* competition binding assays. The effect of **RUPB-61** on the binding of a target kinase to its test probe is reported as a percentage of enzyme binding in the presence of vehicle alone. Blue circles represent the major human off-targets, CIT, ROCK1 and ROCK2. TKL: Tyrosine Kinase-like; STE: Ste kinases; CK1: Casein Kinase 1; AGC: Protein Kinase A, G and C; CAMK: calmodulin kinase; CMGC: Cyclin-dependent kinases, Mitogen-activated protein kinases, Glycogen synthase kinases and CDC-like kinases.

**Supplementary Figure 5: Experimental structure of RUPB-60 bound to PvPKG.** Compared to the structure with **RUPB-61**, the methyl substituent on the **RUPB-60** (green) piperidine evidently leads to the resolution of Ala816 (grey) in the C-term tail (dotted line) (Ala823 in PfPKG).

**Supplementary Figure 6: FEP+ correlation plot comparing prediction to experiment**. A congeneric set of 35 ligands was used in the **(A)** IFD-MD model of PfPKG **(B)** experimental PvPKG structure.

## Supplementary Tables

**Supplementary Table 1: Natural variations in PfPKG open reading frame.** Bioinformatics analyses of PfPKG genomic sequence in MalariaGen isolates.

**Supplementary Table 2: *In vivo* ADMET properties of RUPB-61.** PK properties in mice after oral and intravenous dosing of 10 mg/kg compound. t_1/2_: half-life; T_max_: maximum time; C_max_: maximum concentration; AUC: area under the concentration-time curve; AUC_last_: AUC last observed concentration; AUC_INF__obs: AUC extrapolated to infinity, based on the last observed concentration; CI__obs_: Observed total plasma clearance.

**Supplementary Table 3: Crystallography data collection and refinement.** X-ray structures of **RUPB-60** and **RUPB-61** bound to *P. vivax* PKG were solved and deposited in the PDB database.

**Supplementary Table 4: Individual predicted ΔG from the IFD-MD PfPKG model for a congeneric set of 35 ligands versus their experimental ΔG. \***Experimental ΔG was inferred from IC_50_ values using the relationship IC_50_ ∼ Kd and ΔG = -RTLn(Kd).

## Notes

### Competing Interest Statement

The authors have declared no competing interest.

### Summary of Updates

Results section updated with new data; Discussion revised.

