## Supplementary Information for "An orally available PfPKG inhibitor blocks sporozoite infection of the liver"

### Supplementary Figure 1

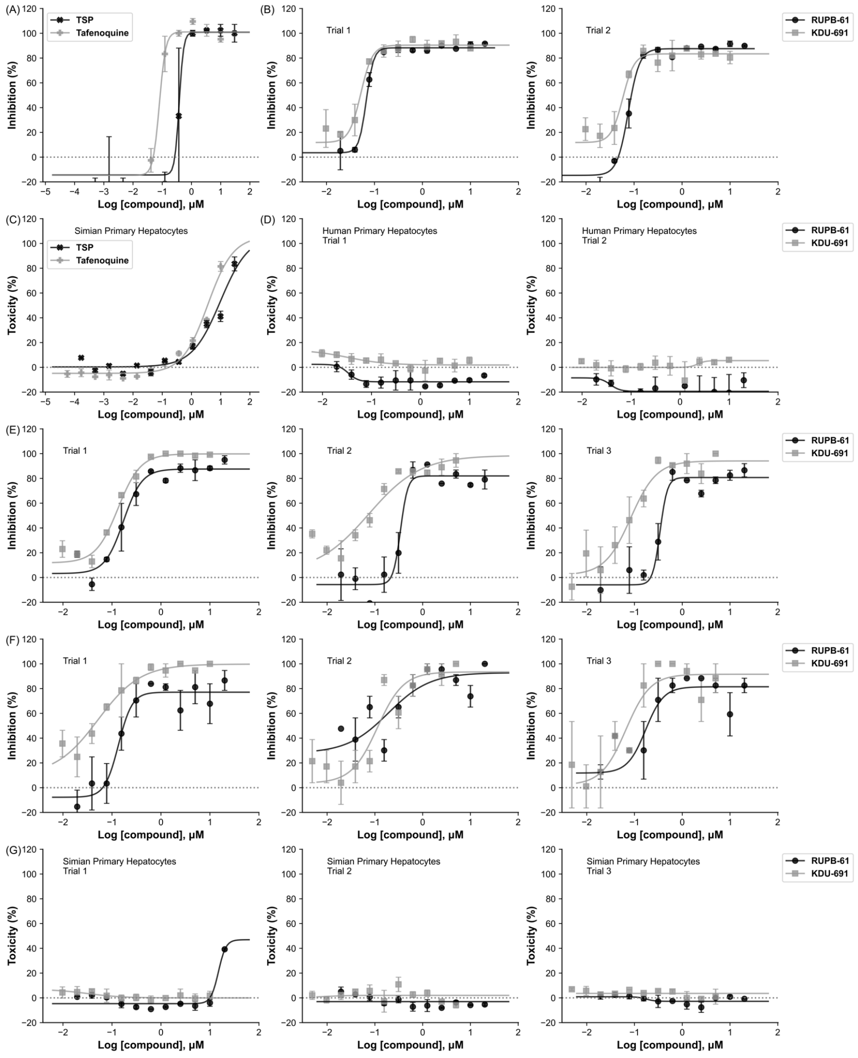

Supplementary Figure 2

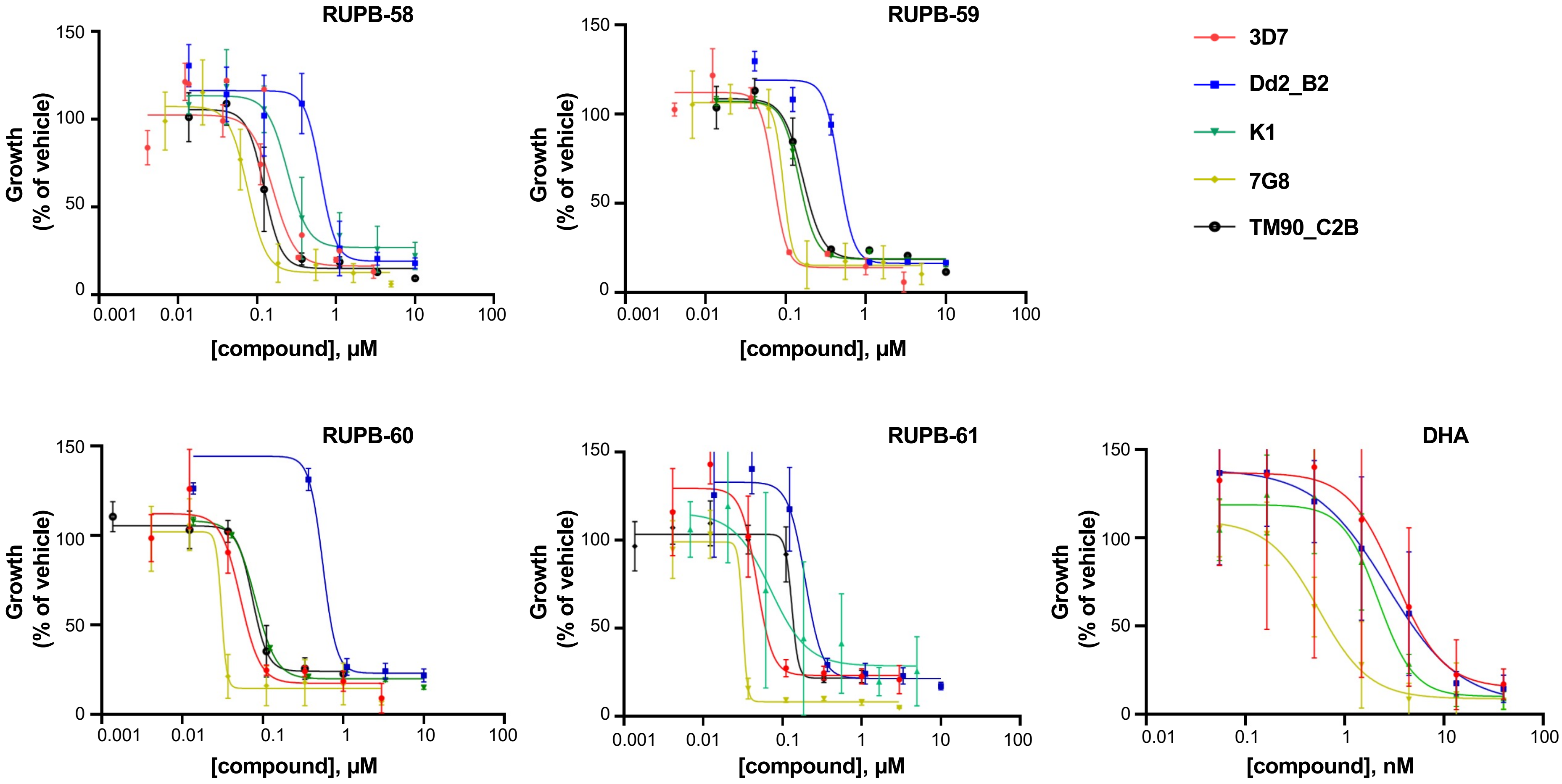

**Supplementary Figure 3**

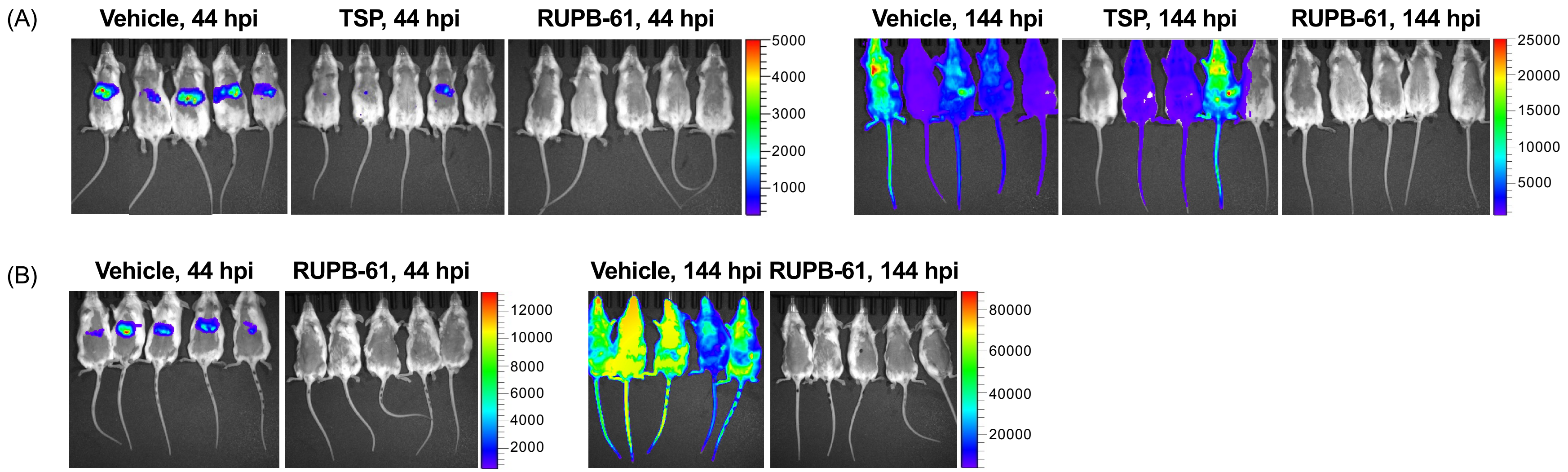

**Supplementary Figure 4**

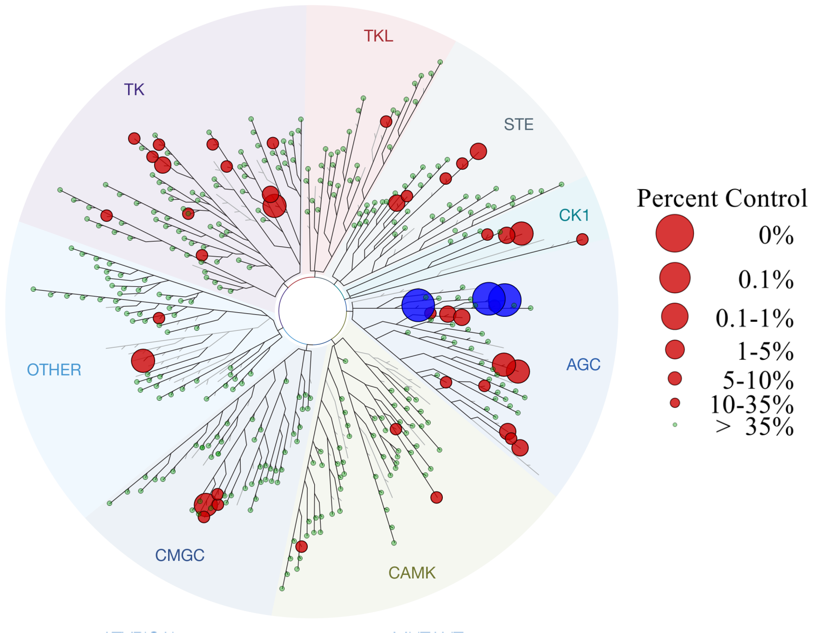

Supplementary Figure 5

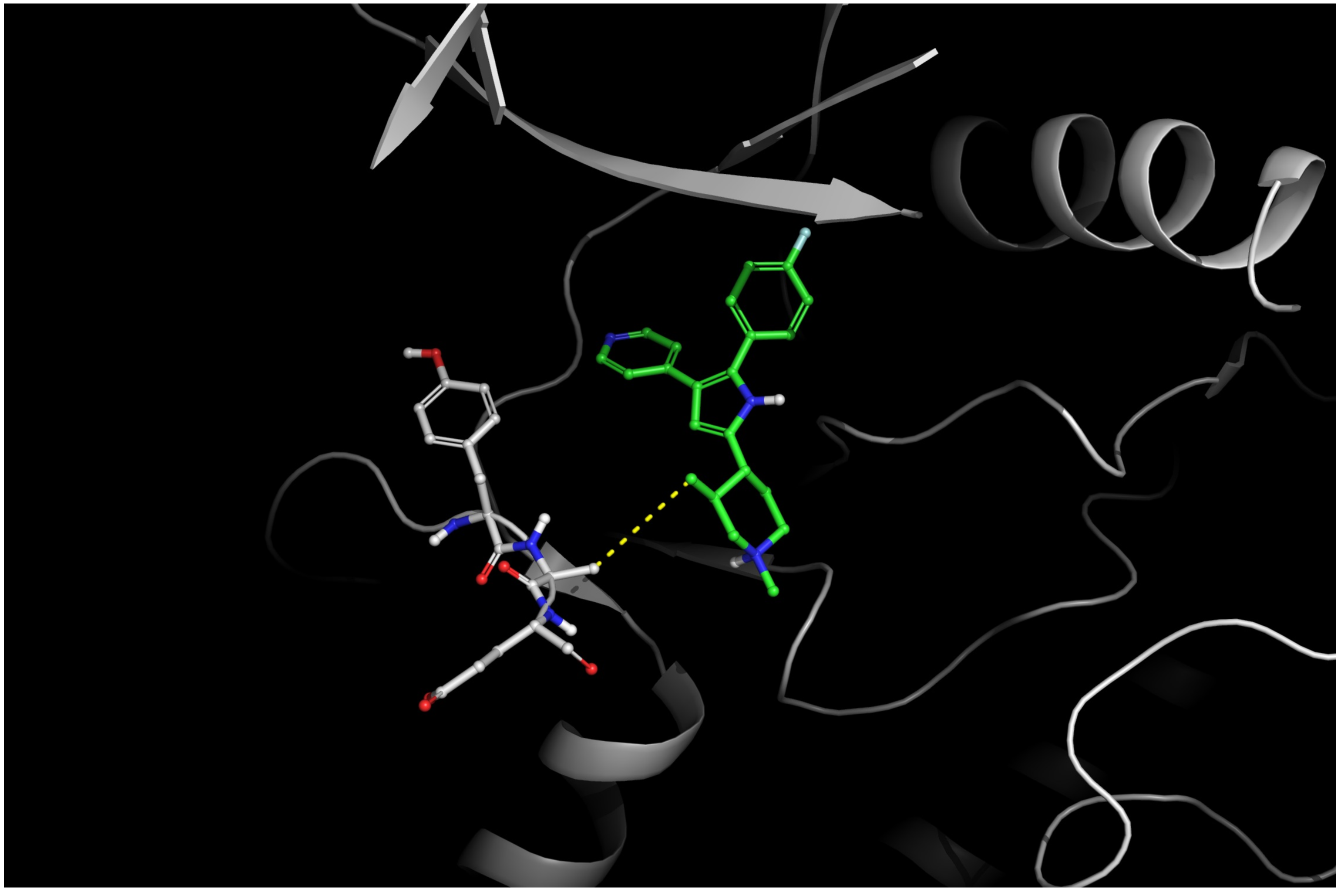

**Supplementary Figure 6**

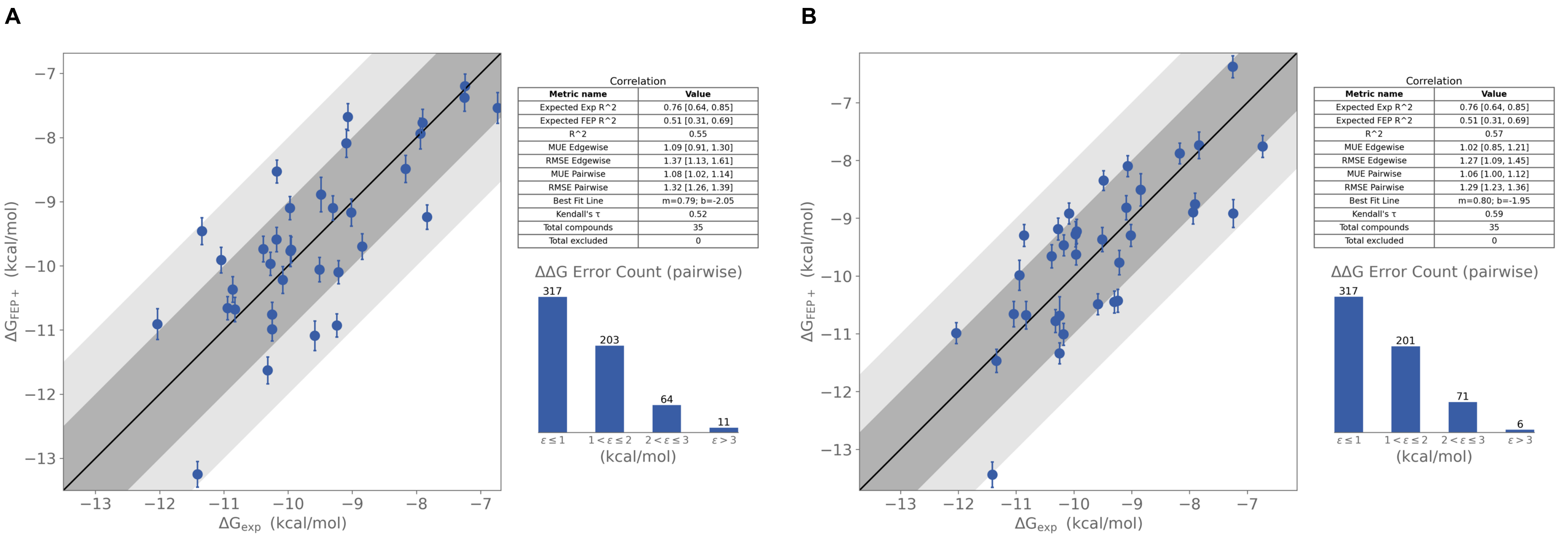

**Supplementary Table 1**

https://nam02.safelinks.protection.outlook.com/?url=https%3A%2F%2F1drv.ms%2Fx%2Fc%2F7ca408f3df9d56b3%2FEVL4-MwEPN5EilKekKIEHB8BpQfu87tHA_0dxwIgl7cNEA%3Fe%3DCm5v5C&data=05%7C02%7Cbhanotpu%40njms.rutgers.edu%7C6ddd7f42f99c4a35e93908ddba736141%7Cb92d2b234d35447093ff69aca6632ffe%7C1%7C0%7C638871726889769208%7CUnknown%7CTWFpbGZsb3d8eyJFbXB0eU1hcGkiOnRydWUsIlYiOiIwLjAuMDAwMCIsIlAiOiJXaW4zMiIsIkFOIjoiTWFpbCIsIldUIjoyfQ%3D%3D%7C0%7C%7C%7C&sdata=qjv6gDeKjRK5gTcei5dIjSeG4MSn1NbHqCcF48H%2FKt0%3D&reserved=0

**Supplementary Table 2**

| PO | $T_{1/2}$ (hr) | $T_{max}$ (hr) | $C_{max}$ (ng/ml) | $C_{max}$ (μM) | $AUC_{last}$ (hr*ng/ml) | $AUC_{last}$ (μM*hr) | $AUC_{INF\_obs}$ (hr*ng/ml) | AUC_%Extrap_obs (%) | Cl_obs (ml/min/kg) | Bioavailability %F |
|---|---|---|---|---|---|---|---|---|---|---|
| Mouse 1 | 8.97 | 6.00 | 170.00 | 0.49 | 2121.20 | 6.07 | 2623.41 | 19.14 | 63.53 | |
| Mouse 2 | 9.71 | 6.00 | 242.00 | 0.69 | 2612.98 | 7.48 | 3370.88 | 22.48 | 49.44 | |
| Mouse 3 | 7.46 | 0.50 | 248.00 | 0.71 | 2906.46 | 8.32 | 3335.65 | 12.87 | 49.97 | |
| Average | 8.71 | 4.17 | 220.00 | 0.63 | 2546.88 | 7.29 | 3109.98 | 18.16 | 54.31 | 91% |

| IV | $T_{1/2}$ (hr) | $T_{max}$ (hr) | $C_{max}$ (ng/ml) | $C_{max}$ (μM) | $AUC_{last}$ (hr*ng/ml) | $AUC_{last}$ (μM*hr) | $AUC_{INF\_obs}$ (hr*ng/ml) | AUC_%Extrap_obs (%) | Cl_obs (ml/min/kg) | $MRT_{INF\_obs}$ hr | Vss_obs L/kg |
|---|---|---|---|---|---|---|---|---|---|---|---|
| mouse 1 | 10.82 | 0.08 | 998.00 | 2.86 | 2400.72 | 6.87 | 3062.87 | 21.62 | 54.42 | 15.22 | 49.69 |
| mouse 2 | 14.92 | 0.08 | 1340.00 | 3.83 | 2698.95 | 7.72 | 3934.67 | 31.41 | 42.36 | 20.35 | 51.71 |
| mouse 3 | 9.59 | 0.08 | 1250.00 | 3.58 | 2687.75 | 7.69 | 3268.89 | 17.78 | 50.99 | 13.23 | 40.47 |
| Average | 11.78 | 0.08 | 1196.00 | 3.42 | 2595.81 | 7.43 | 3422.15 | 23.60 | 49.25 | 16.27 | 47.29 |

#### Supplementary Table 3

| **Sample** | **RUPB-60** | **RUPB-61** |
|---|---|---|
| **PDB Code** | **9P75** | **9P74** |
| **Data Collection** | | |
| Unit-cell parameters (Å, °) | a=190.52<br>b=117.85<br>c=67.31<br>b=94.0 | a=190.17<br>b=117.35<br>c=67.31<br>b=93.55 |
| Space group | C2 | P432 |
| Resolution (Å)[1] | 47.51-2.90<br>(3.06-2.90) | 49.91-2.80<br>(2.94-2.80) |
| Wavelength (Å) | 0.9786 | 0.9786 |
| Temperature (K) | 100 | 100 |
| Observed reflections | 209,792 | 164,453 |
| Unique reflections | 32,911 | 34,947 |
| <I/s(I)>[1] | 10.1 (2.0) | 10.9 (1.9) |
| Completeness (%)[1] | 99.9 (99.9) | 96.1 (100) |
| Multiplicity[1] | 6.4 (6.8) | 4.7 (5.0) |
| $R_{merge}$ (%)[1, 2] | 11.0 (0.912) | 8.9 (94.2) |
| $R_{meas}$ (%)[1, 4] | 12.0 (98.8) | 10.1 (105.1) |
| $R_{pim}$ (%)[1, 4] | 4.7 (37.6) | 4.5 (46.1) |
| $CC_{1/2}$ [1, 5] | 0.998 (0.862) | 0.998 (0.777) |
| **Refinement** | | |
| Resolution (Å) [1] | 47.41-2.90 | 49.91-2.80 |
| Reflections (working/test)[1] | 31,167/1,711 | 33,283/1,646 |
| $R_{factor}$ / $R_{free}$ (%)[1,3] | 22.1/26.6 | 23.3/27.1 |
| No. of atoms (Protein/ Ligand) | 5,946/26 | 5,940/26 |
| **Model Quality** | | |
| R.m.s deviations | | |
| Bond lengths (Å) | 0.002 | 0.004 |
| Bond angles (°) | 0.460 | 0.488 |
| Mean B-factor (Å$^2$) | | |
| All Atoms | 100.5 | 94.9 |
| Protein | 100.5 | 94.9 |
| Ligand | 106.7 | 109.9 |
| Coordinate error (maximum likelihood) (Å) | 0.44 | 0.43 |
| Ramachandran Plot | | |
| Most favored (%) | 98.5 | 97.7 |
| Additionally allowed (%) | 1.5 | 2.3 |

#### Supplementary Table 4

| Ligand | Predicted Binding (ΔG) (kcal/mol) | Experimental Binding* (ΔG) (kcal/mol) |
|---|---|---|
| **RUPB-001** | -9.6 | -10.2 |
| **RUPB-002** | -10.9 | -9.2 |
| **RUPB-005** | -9.2 | -7.8 |
| **RUPB-006** | -7.9 | -7.9 |
| **RUPB-008** | -8.7 | -10.2 |
| **RUPB-009** | -11.1 | -9.6 |
| **RUPB-012** | -7.5 | -6.7 |
| **RUPB-013** | -7.2 | -7.2 |
| **RUPB-014** | -8.9 | -9.5 |
| **RUPB-015** | -10 | -9.5 |
| **RUPB-016** | -8.5 | -8.2 |
| **RUPB-017** | -9.2 | -9 |
| **RUPB-018** | -10.4 | -10.9 |
| **RUPB-019** | -7.5 | -7.3 |
| **RUPB-021** | -7.6 | -9.1 |
| **RUPB-024** | -7.7 | -7.9 |
| **RUPB-028** | -9.8 | -10 |
| **RUPB-029** | -10.2 | -10.1 |
| **RUPB-030** | -9.7 | -10 |
| **RUPB-031** | -10.6 | -10.9 |
| **RUPB-032** | -11.6 | -10.3 |
| **RUPB-033** | -9.7 | -8.8 |
| **RUPB-034** | -10 | -10.3 |
| **RUPB-035** | -10.1 | -9.2 |
| **RUPB-036** | -9.1 | -10 |
| **RUPB-037** | -9.1 | -9.3 |
| **RUPB-59** | -10.7 | -10.8 |
| **RUPB-61** | -9.9 | -11 |
| **RUPB-63** | -9.3 | -11.3 |
| **RUPB-65** | -13.2 | -11.4 |
| **RUPB-68 R** | -10.7 | -10.2 |
| **RUPB-68 S** | -11.1 | -10.2 |
| **RUPB-69** | -10.9 | -12 |
| **RUPB-72** | -11.1 | -10.7 |
| **RUPB-73** | -8.1 | -9.1 |
| **TSP** | -9.7 | -10.4 |
